# Mapping oak wilt disease using phenological observations from space

**DOI:** 10.1101/2023.05.25.542318

**Authors:** J. Antonio Guzmán Q., Jesús N. Pinto-Ledezma, David Frantz, Philip A. Townsend, Jennifer Juzwik, Jeannine Cavender-Bares

## Abstract

Protecting the future of forests relies on our ability to observe changes in forest health. Thus, developing tools for sensing diseases in a timely fashion is critical for managing threats at broad scales. Oak wilt —a disease caused by a pathogenic fungus (*Bretziella fagacearum*)— is threatening oaks, killing thousands yearly while negatively impacting the ecosystem services they provide. Here we propose a novel workflow for mapping oak wilt by targeting temporal disease progression through symptoms using land surface phenology (LSP) from spaceborne observations. By doing so, we hypothesize that phenological changes in pigments and photosynthetic activity of trees affected by oak wilt can be tracked using LSP metrics derived from the Chlorophyll/Carotenoid Index (CCI). We used dense time-series observations from Sentinel-2 to create Analysis Ready Data across Minnesota and Wisconsin and to derive three LSP metrics: the value of CCI at the start and end of the growing season, and the coefficient of variation of the CCI during the growing season. We integrate high-resolution airborne imagery in multiple locations to select pixels (*n =* 3,872) from the most common oak tree health conditions: healthy, symptomatic for oak wilt, and dead. These pixels were used to train an iterative Partial Least Square Discriminant (PLSD) model and derive the probability of an oak tree (i.e., pixel) in one of these conditions and the associated uncertainty. We assessed these models spatially and temporally on testing datasets revealing that it is feasible to discriminate among the three health conditions with overall accuracy between 80-82%. Within conditions, our models suggest that spatial variations among three CCI-derived LSP metrics can predict healthily (Area Under the Curve (AUC) = 0.98), symptomatic (AUC = 0.89), and dead (AUC = 0.94) oak trees with low false positive rates. The model performance was robust across different years as well. The predictive maps were used to guide local stakeholders in locating disease hotspots for ground verification and subsequent decision-making for treatment. Our results highlight the capabilities of LSP metrics from dense spaceborne observations to map diseases and their importance for monitoring changes in biodiversity at large scales.

## 1. Introduction

The oak (*Quercus*) lineage is one of the most important groups of trees in North American temperate forests. This tree lineage comprises 172 oak species in the United States and Mexico, contributing to more than 20% of total aboveground biomass (Cavender-Bares, 2019). As a result, oak trees play an invaluable role in the biodiversity, structure, and ecosystem functioning of temperate forests, in part as a consequence of their high abundance and diversity. In the U.S., oaks contribute nearly $22.3 B annually in net ecosystem services from wood products to climate protection, and air quality regulation (Cavender-Bares et al., 2022). Alarmingly, oak trees are facing major threats from diseases and pests that stand to impact their role in ecosystem functioning and the provision of ecosystem services. Protecting the future of oak forests relies on our ability to observe changes in their health conditions. Early detection followed by rapid response gives the highest probability of disease management success. Thus, developing tools for remotely sensing forest health in a timely fashion is critical for managing threats at a large scale.

One of the most lethal oak tree diseases, particularly of red oak species, is oak wilt (Gibbs and French, 1980). Oak wilt is caused by an invasive fungal pathogen *Bretziella fagacearum* (Bretz, 1952; de Beer et al., 2017; Hunt, 1956). Spores of the oak wilt fungus are translocated in the sapstream of the vascular system of infected oak trees. In response, infected trees produce tyloses to isolate or slow lateral or vertical spread through the vessels (Struckmeyer et al., 1954; Yadeta and Thomma, 2013). The efficacy of tylose formation in limiting within tree spread of the pathogen is strongly linked to the vessel diameter and rapidity of the host response, which differs by oak lineage. The white oaks (*Quercus* section *Quercus*) tend to have denser wood with smaller diameter vessels than red oaks (Cavender-Bares and Holbrook, 2001) in which tyloses are more effective at slowing the vascular spread of oak wilt, whereas red oaks (*Quercus* section *Lobatae*) have larger vessels and their immune responses are relatively ineffective in halting the fungus spread (Juzwik et al., 2011; Yadeta and Thomma, 2013). Metabolites produced by the fungus also contribute to the plugging of xylem vessels. Once an oak tree is infected, the pathogen can spread locally belowground or long-distance by above-ground means (Gibbs and French, 1980; Juzwik et al., 2011). The below-ground spread occurs by fungal movement through networks of grafted roots of neighboring trees forming “pockets” or centers of diseased and dead oak trees (Kuntz and Riker, 1950). Above-ground spread is facilitated by insect vectors (Gibbs and French, 1980) such as several species of Nitidulid beetles (Coleoptera: Nitidulidae) that acquire *B. fagacearum* spores when visiting oak wilt sporulating mats on diseased trees and subsequently transmit the spores to fresh xylem-penetrating wounds on healthy oaks (Juzwik et al., 2011). Accurate and early detection of oak wilt facilitates successful treatment of the disease. For example, severing roots around the pockets and removing diseased oaks is one proven tactic that is effective in halting the propagation of oak wilt (Juzwik et al., 2011; Koch et al., 2010). Despite this, the detection of pockets or isolated individual diseased trees is laborious and expensive, particularly at regional scales. Thus, the development of remote sensing tools for detecting diseased trees at large spatial scales is crucial first step for guiding management efforts.

Oak trees in advanced stages of the disease tend to express drought-like symptoms associated with reductions in water content and concentration of green pigments in their leaves (Fallon et al., 2020). Such symptoms lead to spectral signatures that can be detected at the leaf (Fallon et al., 2020) or canopy level using hyperspectral sensors (Sapes et al., 2022). Such symptoms can also be differentiated using three-dimensional color space models (Monahan et al., 2022). However, it remains unclear the extent to which detection is feasible at the spaceborne level where extensive spatial coverage and frequent observations could be highly informative. Although pixel size from spaceborne observations could limit the ability to detect individual trees in comparison with traditional aerial surveys, frequent satellite observations are likely to track the progression of crown wilting in mid-summer, which is considered a key symptom for diagnosing the disease in the Upper Midwest (Haugen et al., 2022). We thus expect the use of phenological metrics that can be derived from these observations to be useful in observing the symptoms of disease progression and mapping the spread of the disease in oak trees at the regional scale.

Here we evaluate the use of the temporal spaceborne observations for the mapping of oak wilt in forests of the Upper Midwest U.S. We present a reproducible, scalable, and open-source workflow to map oak wilt and its impacts across Minnesota and Wisconsin — but applicable to other regions—to advance guidance of disease management efforts by forest managers and stakeholders. In doing so, we hypothesize that phenological changes in pigments and photosynthetic activity of oak trees due to oak wilt can be tracked using phenological metrics derived from the Chlorophyll/Carotenoid Index (CCI) (Gamon et al., 2016) from satellite observations. We evaluate this approach within the spatial and temporal context in conjunction with high-resolution airborne observations of symptomatic trees in pockets of disease from Central Minnesota. This study highlights the use of phenological base metrics from frequent and continuous spaceborne observations for mapping diseases at a large scale to further understand the impacts of tree pathogens on the ecosystem functioning and services in the North American temperate forest.

## 2. Phenological observations as a proxy for detecting oak wilt

Susceptible red oak trees infected with the oak wilt pathogen usually present a rapid wilting of their crown in a period of weeks. Infected white oaks, on the other hand, tend to present scattered wilting of their crown over several to many years (Haugen et al., 2022). The wilting process can vary in timing but frequently occurs in mid to late summer. While a definitive diagnosis of oak wilt disease requires laboratory isolation of the fungus or a DNA test, rapid wilting, particularly of red oaks in a cluster formation extending outward from a previously diseased tree (or trees), is considered a key symptom for on-site diagnosis by trained individuals. In this study, we seek to detect and map oak wilt based on knowledge of the disease progression using temporal observations of a spectral index sensitive to oak wilt symptoms.

Consider the variation over time of the CCI spectral index known to be sensitive to oak wilt (Sapes et al., 2022) (Fig. 1a). At the start of the growing season, the CCI signal from a healthy or symptomatic oak tree expresses sharp increases due to leaf flushing that differentiates it from the CCI signal of a dead oak tree (Fig. 1a). Healthy, uninfected oaks exhibit seasonal variation in CCI as a consequence of leaf maturation and seasonal changes in chlorophyll and carotenoid pigment ratios (Gamon et al., 2016). Wilting in a diseased oak tree produces a more pronounced and earlier reduction in CCI than in a healthy tree. As such, the wilting process leads to a reduction in the mean CCI over the course of the growing season and a symptomatic signal close to the senescence period that resembles dead trees. Higher variability (i.e., standard deviation) and lower mean CCI during the growing session led to higher coefficients of variations in symptomatic oak trees in comparison with healthy or dead oak trees. Thus, metrics that describe phenological events of CCI, including the value at the start of the season (VSS), the value at the end of the season (VES), the variability value as the standard deviation of CCI during the growing season (VGV) or the coefficient of variation (VCV), provide a basis to differentiate healthy, symptomatic, and dead oak trees. These metrics can be normalized locally (i.e., z-scores) using the values of healthy vegetation (e.g., mean and standard deviation), to derive trends between conditions (Fig. 1b). The differentiation of symptomatic, dead, and healthy oak trees ultimately leads to the mapping of the disease.

**Fig 1.**
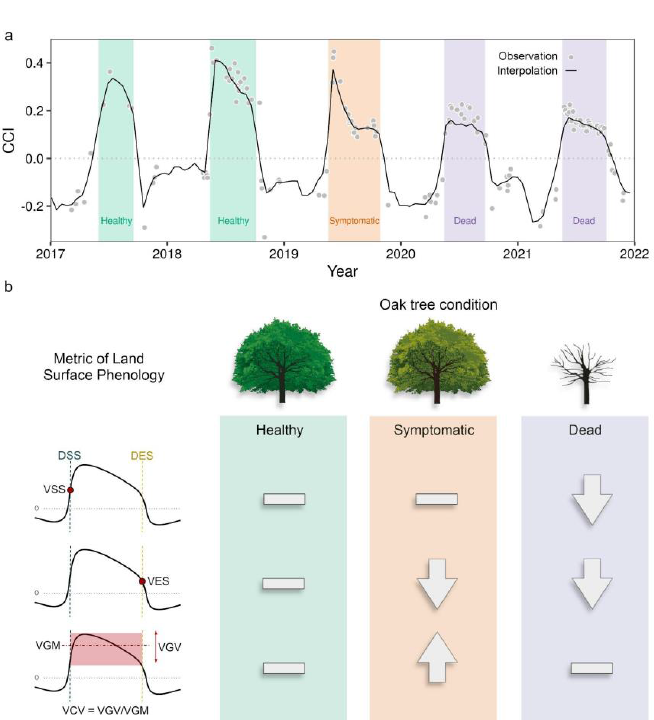
Temporal changes in the Chlorophyll/Carotenoid Index (CCI) of a pixel from an oak tree that died from oak wilt disease (a) and the expected behavior of land surface phenology metrics when oak tree conditions are compared with the surrounding healthy vegetation for a given phenological year (b). Individual points represent satellite observations, while the solid line shows the 16-day interpolation. The vertical color bars represent the evaluation of the three oak tree conditions (green: healthy; orange: symptomatic; purple: dead). The acronyms in panel b represent the day of the start and the day of the end of the green season (DSS and DES), the values at the start and end of the growing season (VSS and VES), the mean and standard deviation value during the growing season value (VGM and VGV), and the value of the coefficient of variation during the green season (VCV). Arrows indicate expected values higher or lower than healthy oak trees, while a dash indicates a value similar to healthy trees.

## 3. Materials and Methods

### 3.1 Study area

Our study site encompasses 365,700 km^2^ in the states of Minnesota (MN) and Wisconsin (WI), U.S.A. These neighboring states present two forest ecological provinces dominated by the Laurentian Mixed Forest and the Eastern Broadleaf Forest (Bailey, 1998). Also present is the Prairie Parkland Province, dominated mainly by grasslands and agriculture (Bailey, 1998), with limited tree cover and a low abundance of oaks and oak wilt. Forest cover is the most dominant vegetation type in both states, with forest cover close to 50% and 48% of the total area in MN (Miles et al., 2016) and WI (Wisconsin Department of Natural Resources, 2020), respectively. Both states present a similar gradient of agricultural cover in the southernmost regions to forested cover in the northern regions. This region is well-suited for testing phenological metrics for oak wilt detection due to the wide distribution of oaks, the current spread of oak wilt, and the strong climatic seasonality with the potential to provide valuable information to forest managers.

### 3.2 Satellite data processing

We used satellite observations from Landsat 8 collection 2 Tier 1 L1TP data (USGS, 2021) and Sentinel-2 A/B Level 1C (Drusch et al., 2012) with a cloud cover of up to 70%. Images were processed through FORCE (version 3.7.7; Frantz 2019); a single computing environment to efficiently process, analyze, and stack satellite observations in a datacube framework. These images were corrected for geometric and radiometric effects, reproject to the USA Contiguous Lambert projection (EPSG:102004), and store as 30 km × 30 km datacube tiles (**Fig. S1**) to create Analysis Ready Data (ARD) (Fig. 2) (i.e., Level 2 ARD). Geometric and radiometric corrections include corrections for atmospheric, topographic, adjacency, and bidirectional reflectance distribution function (BRDF) effects (Buchner et al., 2020; Frantz et al., 2016a; Roy et al., 2017). The creation of ARD also includes masking clouds based on Fmask (Frantz et al., 2018; Zhu et al., 2015; Zhu and Woodcock, 2012). Topographic corrections and enhancements for cloud detection were applied based on a digital elevation model from Copernicus GLO-90 Digital Surface Model accessed through OpenTopography (2021). Landsat 8 scenes from 2017 to 2021 were first ingested as a baseline for co-registration to create monthly multi-annual Landsat 8 near-infrared scenes. These were used to co-register Sentinel-2 to ensure the best possible geometric alignment between scenes and among sensors (Rufin et al., 2021). Then, Sentinel-2 scenes from November 2016 to February 2023 were ingested and processed using FORCE. All the Sentinel-2 bands were downscaled to a 10 m spatial resolution using the ImproPhe algorithm (Frantz et al., 2016b). **Fig. S2** illustrates the total amount of clear sky observations for the years of interest.

**Fig 2.**
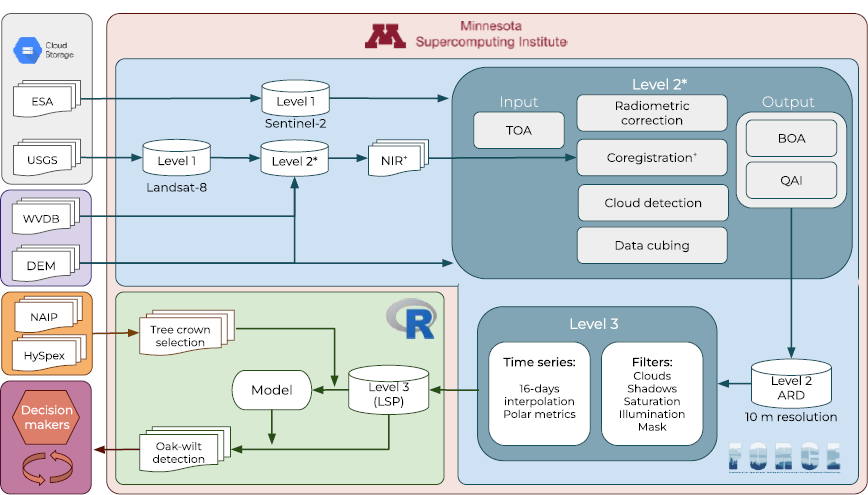
Workflow for acquiring, processing, and analyzing Landsat 8 and Sentinel-2 imagery for retrieving land surface phenology metrics and mapping oak wilt disease.

### 3.3 Time series of Chlorophyll/Carotenoid Index

#### 3.3.1 Chlorophyll/Carotenoid Index

Using Sentinel-2 ARD, we calculated the chlorophyll/carotenoid index (CCI) (Gamon et al., 2016) to target physiological responses sensitive to wilting. Precisely, the CCI index was estimated as:

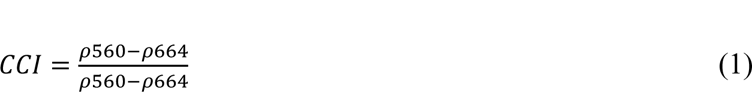

where *p*560 and *p*664 are bands 3 and 4 of Sentinel-2. Originally this index was developed to track the evergreen forest phenology using the Moderate Resolution Imaging Spectroradiometer (MODIS) (Gamon et al., 2016), but it has also been found to be sensitive to mixed deciduous forest phenology (Wong et al., 2020) and oak wilt (Sapes et al., 2022). Although other indices are sensitive to oak wilt symptoms (e.g., Sapes et al., 2022), we use CCI because of its relationship to physiological stresses and strong correlation/redundancy with other indices. The kernel density of Normalize Difference of Vegetation Index (kNDVI) (Camps-Valls et al., 2021) was also calculated to be used as a forest mask in the following procedures. The CCI and kNDVI estimation, the temporal interpolation of observations, and the derivation of phenological metrics were also conducted in FORCE using the ‘force-higher-level’ chain processing submodule as detailed below.

#### 3.3.2 Land surface phenology

Our approach to using LSP metrics to detect oak wilt is based on the premise that visible oak wilt symptoms are not likely to be expressed during the early growing season (e.g., when the transmission tends to occur), but in the late-growing season after a period of incubation. We test the extent to which LSP metrics using spectral indices related to wilting symptoms can discriminate between symptomatic and dead trees (early growing season) and symptomatic and healthy oak trees (late growing season).

LSP metrics are calculated on a per-pixel basis to create annual time series observations following Brooks et al. (2020) and Frantz et al., (2022). Poor-quality pixels flagged as cloud, cloud shadow, or snow as well as saturated and sub-zero reflectance pixels are excluded from the analysis. The CCI was interpolated to a 16-day observation period using an ensemble of Radial Basis Function convolution filters as described by Schwieder et al., (2016) and Frantz et al., (2022). This filter is based on retrieving a weight value computed from a Gaussian distribution function on a set of observations retrieved from a given kernel width (i.e., days windows) at a specific period (detail in Frantz et al., 2022). We used three kernel widths of 8, 16, and 32 days and a kernel cutoff value that preserves 95% of the area under the Gaussian curve as employed by Frantz et al. (2022), which gives preference to kernel widths with a higher number of observations for the final estimation. No-data values are likely to occur during winter when there are few clear-sky satellite observations, but a linear interpolation is then performed to fill the remaining data gaps which are likely to be negative values and considered as zero for the estimation of the LSP.

LST metrics were computed for each year between 2017 and 2022 using a polar transformation-based method (Brooks et al., 2020; Frantz et al., 2022). Days of the year (DOY) are converted to radians and CCI is transformed into a cartesian space using angles. An index value from a given observation period will thus result in a transformed value with a [*x*, *y*] cartesian coordinate. From all the cartesian coordinates a long-term polar average vector is computed to describe the central tendency of the annual peak of the phenological year. The diametric opposite angle of this average vector is then estimated to characterize the direction to the lowest values and thus, the long-term start of the phenological year. The timing of the peak and the start of the phenological year can vary from pixel to pixel and gradually change according to its latitude. The long-term start of the phenological year is then used to divide the whole interpolated time series into annual slices of phenological years. The start of the phenological year is then re-estimated on each annual slice to consider the potential inter-annual variability following Frantz et al. (2022). This fine-tuning of the phenological year consists in re-computing the polar average vector and its diametric opposite angle per season to further describe phenological years with dynamics starting and ending days.

Once the dynamic phenological years are defined, values from spectral indices were converted from zero to one in cumulative proportions which were used to characterize cumulative milestones for the starting and ending of growing seasons within a phenological year based on thresholds. We chose phenological thresholds at 0.15 and 0.85 of the cumulative proportions to define the beginning and end of the growing season (e.g., green season). Using these thresholds, five LSP metrics were derived from CCI (as explained in *Section 2*): VSS, VES, VGM, VGV, and VCV as VGV/VGM. VSS and VES are defined as the CCI value at 0.15 and 0.85 of the cumulative threshold, while VGM and VGV as the mean and standard deviation of the CCI between the start and end of the growing season. The previous LSP metrics except for VCV were directly computed in FORCE. In addition, the value of the peak of the growing season (VPS) was derived from kNDVI and used as a forest mask if it exceeds a threshold of 0.5.

We normalized the LSP metrics derived from CCI (i.e., VSS, VSS, and VCV) each year by tile based on z-scores to further reduce the inter-annual and spatial variability of the phenological observations. We used the annual VSS to identify healthy forest cover and then used VSS values higher than 0.25 to mask forested areas given their clear differentiation from other cover types. The mean and standard deviation of the resulting masked pixels were used to compute the z-scores (e.g., (LSP metric - LSP mean) / LSP standard deviation). The LSP metrics estimated are available at https://app.globus.org/file-manager?origin_id=d5f9b461-7d6e-442b-87ed-be8aa2ca6763&path=%2F.

### 3.4 Detection of oak wilt and accuracy assessment

#### 3.4.1 Oak wilt dataset

We developed datasets of observations of symptomatic trees from three years of airborne imagery to guide the selection of pixels to model oak wilt presence from metrics of LSP. Aerial data included 0.3 m multispectral National Agriculture Imagery Program (NAIP) scenes acquired between July 15^th^ to August 15^th^, 2019 across eight tiles of ARD (900 km^2^ each) and 1 m resolution HySpex VNIR-1800 (NEO, Oslo, Norway) scenes captured in two surveys over six locations on July 24^th^ 2018, and for one location on July 24^th^ 2021. The HySpex images were processed following Liu et al. (2021) and Queally et al. (2022), and we used composite images of 664, 562, and 491 nm to create true color images to identify oak wilt locations. All aerial imagery was co-registered to the average red band of Sentinel-2 ARD for the concurrent period using AROSICS (Scheffler et al., 2017).

Once the airborne imagery was co-registered, we then proceeded with the digitalization of points. This was done with the guide of digitized polygons of pockets of disease created by the Department of Natural Resources (DNR) of Minnesota (Department of Natural Resources, 2021). These disease pockets are ground-checked areas based on 2019 National Agriculture Imagery Program (NAIP) imagery, reports from DNR foresters and the U.S. Department of Agriculture Forest Service, and windshield surveys (Department of Natural Resources, 2021). These polygons were distributed over several counties across Minnesota, and close to or within them it is possible to visualize trees with oak wilt symptoms, which we infer are mostly red oaks species (*Quercus rubra* or *Quercus ellipsoidalis*). These polygons were manually co-registered to our 2019 NAIP co-registered imagery and used as a guide for selecting pixels (i.e., trees) in areas affected by the disease. We manually digitized points on tree crowns that appeared to be symptomatic of oak wilt (i.e., wilted), healthy, or dead within or around the disease pockets on the high-resolution airborne imagery. We assumed that if an oak crown looked symptomatic, was within or close to a disease pocket, and appeared to be dead in subsequent years, it is likely that the tree was killed by oak wilt. A single point was digitized on symptomatic or dead crown within or close to small disease pockets (∼ 0.5 ha), and between two and three points within large pockets to avoid potential spatial autocorrelation among pixels. All trees classified as symptomatic trees in 2018 and 2019 were assessed for mortality using NAIP imagery of the following years. In total, 3,872 points were digitized (**Table S1** and **Fig. S3**). We then used these points to extract pixels for the LSP metrics (*Section 2.3.3*) and to develop the models for detecting the disease (*Section 2.4.2*). The digitalization of points was performed using QGIS 3.22 (QGIS Development Team, 2022) with a USA Contiguous Lambert projection (EPSG:102004) and a fixed window scale of 1:1000.

#### 3.4.2 Model development and assessment

We built machine learning models to detect oak wilt from the LSP metrics, with the ultimate objective of a map of the probability of oak wilt presence. We randomly split 60% of samples (i.e., pixels) from the 2019 dataset to train the classification algorithms. Specifically, the data splitting procedure was conducted on samples from each tile based on their condition (i.e., symptomatic, healthy, or dead) to ensure a balanced representation of conditions by spatial locations during the algorithm’s training. The training dataset encompassed 1,984 samples (661 samples per condition) at the end of this procedure. We used the remaining 40% of samples (*n =* 1,520) to test the performance of classification at the spatial level as well as at the temporal using the 2018 and 2021 datasets (*n* = 370).

Using the established framework, we trained and selected classification algorithms. Six widely-used classification algorithms that required little or no tuning parameters for their computation were implemented using the caret package in R (Kuhn, 2008) (Table 1). During training, we avoided potential spatial auto-correlation within a set of scenes by using an 8-fold spatial cross-validation in which ARD tiles comprised the spatial component. We iterated this optimization procedure 100 times using 80% of the training dataset during each iteration using the CAST package of R (Meyer et al., 2023).

**Table 1.** List of classification algorithms and their corresponding tuning parameters used to differentiate oak tree conditions.

| Algorithm | Abbreviation | Function | Library | Turning parameter |
| --- | --- | --- | --- | --- |
| Linear Discriminant Analysis | LDA | lda | MASS (Venables and Ripley, 2002) | — |
| Quadratic discriminant analysis | QDA | qda | MASS (Venables and Ripley, 2002) | — |
| Support Vector Machine with Linear Kernel | SVM | svmLinear | kernlab (Karatzoglou et al., 2004) | Cost = 0.3 |
| k-Nearest Neighbors | KNN | knn | — | k = 33 |
| Partial Least Squares Discrimination | PLSD | pls | pls (Mevik and Wehrens, 2007) | ncomp = 3 |
| Random forest | RF | rf | randomForest (Liaw and Wiener, 2002) | mtry = 1 |

We selected the final model for implementation based on accuracy assessment and kappa statistics. The best-performing model was further evaluated with four main descriptors of performance: i) balance accuracy; ii) the sensitivity, or the true-positive rate, which measures the probability of which a pixel condition is predicted correctly for all samples of that condition; iii) the specificity or 1-the false-positive rate, which measures the probability of which non-event conditions are predicted as non-event conditions, and iv) the F1 which describes the average of precision and recall (Kuhn, 2008). Since our goal is to produce maps of probabilities, we examine the selected model further by estimating the receiver operating characteristic (ROC) curves and their area under the curve (AUC) among the condition. We identified the probability level at which there was a balance between sensitivity and 1 – specificity to inform ‘*cutoff’* values to produce maps. The macro and micro effects of multiclass classification were also estimated using ROC curves to evaluate the performance independently of the conditions or potential imbalance among conditions.

#### 3.4.3. Mapping of oak wilt

We applied the selected predictive model to three annual LSP metrics from 2017 to 2022 to map the probability of the oak tree condition in Minnesota and Wisconsin. We applied 100 iterative models to generate a mean probability value and estimated uncertainty as the amplitude between the upper and lower limits of the confidence intervals at 95% of the predicted probabilities per condition, assuming that these follow a normal distribution. We used the sum of amplitudes among conditions as a descriptor of the overall uncertainty of the model on the presented maps (*Section 4.3*). We showcase areas outside of our training tiles affected by oak wilt disease to prove its potential to guide stakeholders. The predicted maps are available at https://app.globus.org/file-manager?origin_id=2ad70821-cc5a-424e-aa72-8553d2bb45eb&path=%2F.

## 4. Results

### 4.1 Datasets evaluation

The LSP metrics differentiate healthy, symptomatic, and dead oak trees within each phenological year of evaluation (Fig. 3). VSS clearly differentiates pixels from healthy oak crowns from symptomatic and dead crowns, with the highest values in healthy trees, and lowest in dead ones (Fig. 3a). For VES, symptomatic and dead oak trees tend to display similar values, but healthy crowns show higher values (Fig. 3c). Similarly, VCV differentiates symptomatic trees from healthy and dead oak trees because symptomatic crowns tend to have higher values than healthy or dead crowns. (Fig. 3e). The z-score transformation to normalize the LSP variation among tiles enhanced this differentiation and our ability to make accurate inferences of crown health. During the three years of evaluation (Fig. 3b, d, and f), healthy oak trees always have values that tend towards zero, while symptomatic and dead crowns tend to be differentiated by their means (except for VCV in dead oak trees). Differentiation among oak tree conditions using the land surface phenology (LSP) metrics and their transformations are also observed spatially in our 2019 dataset (**Fig. S4**). Maps of the three evaluated LSP metrics across Minnesota and Wisconsin are presented in **Fig. S5.**

**Fig. 3.**
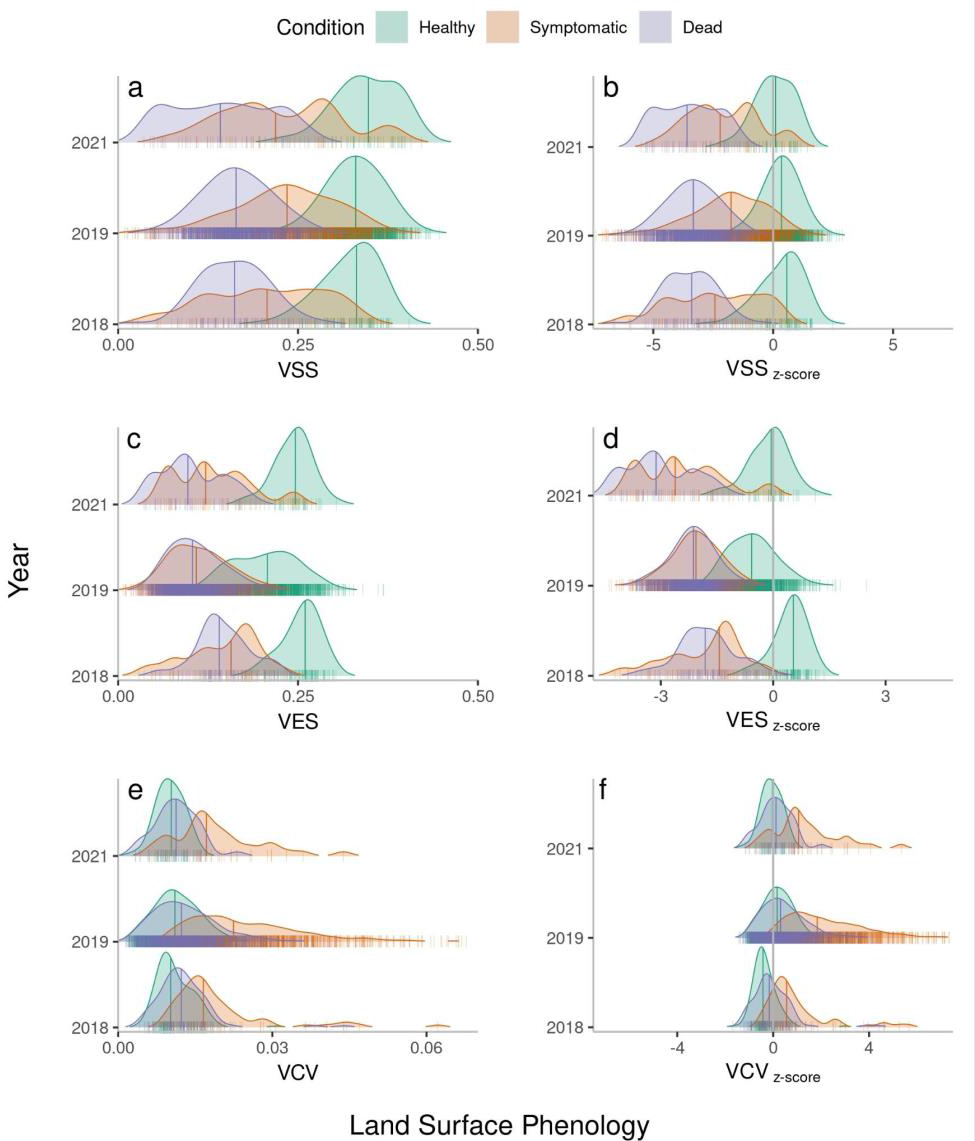
Ridgeline plots for land surface phenology metrics (a, c, e) comparing three oak tree conditions in 2018, 2019, and 2021 and their z-score normalization (b, d, f). Metrics shown are the Value at the Start of the Season (VSS) (a, b), the Value at the end of the Season (VGV) (c, d), and the Value of the Coefficient of Variation (VCV) (e, f). The curves describe the kernel density distributions with different bandwidths for better visualization. Each small vertical line represents a pixel (i.e., an oak tree crown), while the large vertical lines within the density distribution represent the 50^th^ percentile.

### 4.2 Assessment of the predictive model

#### 4.2.1 Assessment of the models

The six machine learning models we tested credibly discriminated pixels among healthy, symptomatic, or dead oak trees (Fig 4). Overall, the mean accuracy for the training dataset ranges between 0.76 and 0.81 among all the classifiers, and between 0.80 and 0.82 for the testing dataset. Similarly, the mean kappa value for training data ranged between 0.64 and 0.72 (0.68 ± 0.01) and between 0.69 and 0.73 (0.71 ± 0.01) for testing data. Among the algorithms we tested, LDA, PLSD, and KNN exhibited the highest mean accuracy close to 0.79 for the training dataset. For the testing datasets, QDA had a mean accuracy of 0.82 closely followed by KNN, SVM, LDA, and PLSD with values close to 0.81. RF gave the lowest classification performance on both training and testing datasets with accuracies of 0.78 and 0.80, respectively. Given that the PLSD performance was among the highest and is readily applied and interpreted in comparison with QDA or SVM, we selected the PLSD model to map symptomatic trees and further test spatial and temporal classification performance.

**Fig 4.**
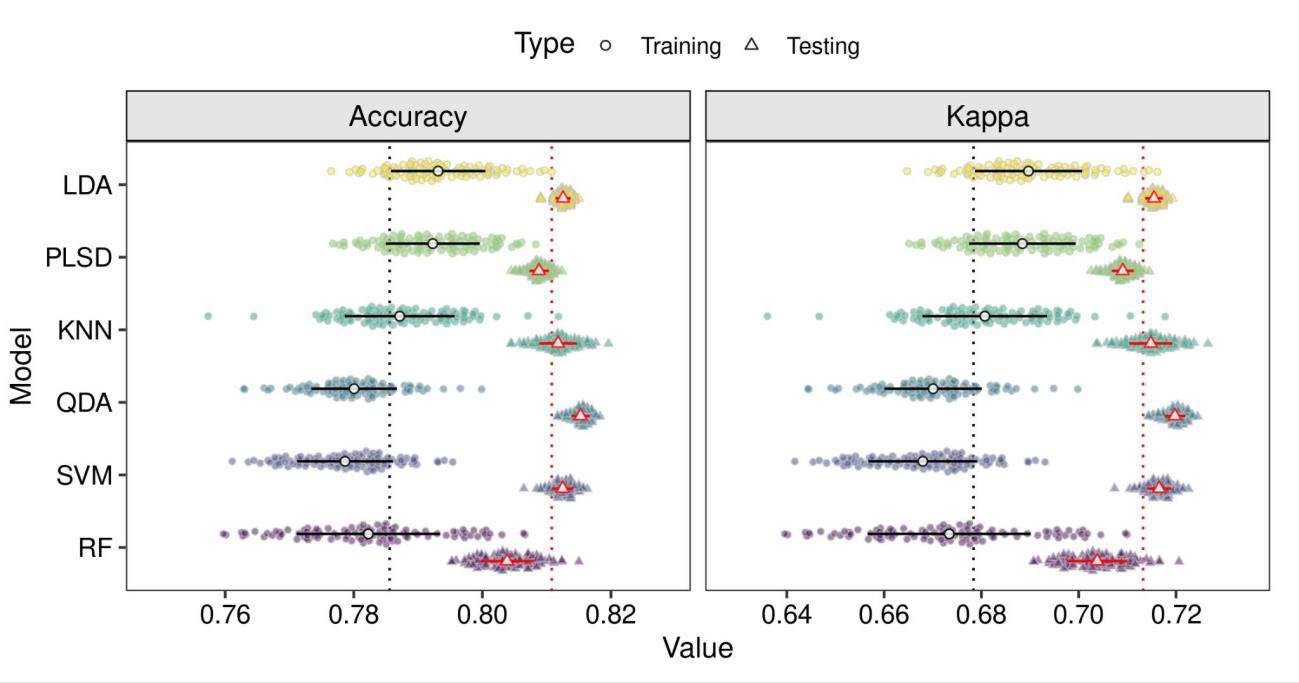
Performance of the classification algorithm for predicting oak tree conditions (i.e., healthy, symptomatic to oak wilt, and dead) on normalized metrics of land surface phenology. Points in a color gradient represent the variability of the repeated model training (*n* = 100). Solid white points represent the mean value, and the error bars show the standard deviation. Vertical dotted lines represent the overall mean of training (black) or testing (red) regardless of the classifiers. The acronyms of the classification algorithms are described in Table 1.

#### 4.2.2 Spatial assessment

Spatial evaluation of the model built using the 2019 testing dataset reveals that the classification performance varies across observation tiles (Fig. 5). The balanced classification accuracy of pixels from healthy oak trees was highest and ranged between 0.83 and 0.95 among tiles; for pixels from dead and symptomatic oak trees it ranged from 0.76 to 0.92, and from 0.67 to 0.88, respectively. Overall, the balanced accuracy of classification for healthy oak trees was higher and less variable (0.90 ± 0.04) among tiles than for dead (0.85 ± 0.04) or symptomatic oak trees (0.80 ± 0.06). We found this same pattern for the sensitivity, specificity, and F1 parameters (Fig. 5). Most of the class confusion was between dead and symptomatic oak trees **(Fig. S6)**. For instance, in tiles where symptomatic oak trees had higher sensitivity or specificity values, dead oak trees showed lower values, and vice versa. Our spatial assessment also indicates that mapping in the three southernmost tiles (i.e., X0016_Y0025, X0017_Y0026, and X0017_Y0027) performed worse than average. However, this performance was always greater than 0.7 of balanced accuracy. In addition, the ROC curves and their AUC demonstrate that across tiles pixels from healthy oak trees are more readily classified (AUC = 0.94 ± 0.05), than dead (AUC = 0.90 ± 0.05) or symptomatic oak trees (AUC = 0.84 ± 0.08) (**Table S2**). These curves also reveal a congruent multiclass classification performance independently of the conditions (i.e., macro) (AUC = 0.89 ± 0.05) or potential class imbalance (i.e., micro) (AUC = 0.88 ± 0.06). The probability of ‘*cutoff*’ from these ROC curves to obtain a balance between sensitivity and 1 – specificity within conditions was higher in healthy oak trees (0.41 ± 0.04) than in dead (0.35 ± 0.02) or symptomatic trees (0.32 ± 0.01).

**Fig 5.**
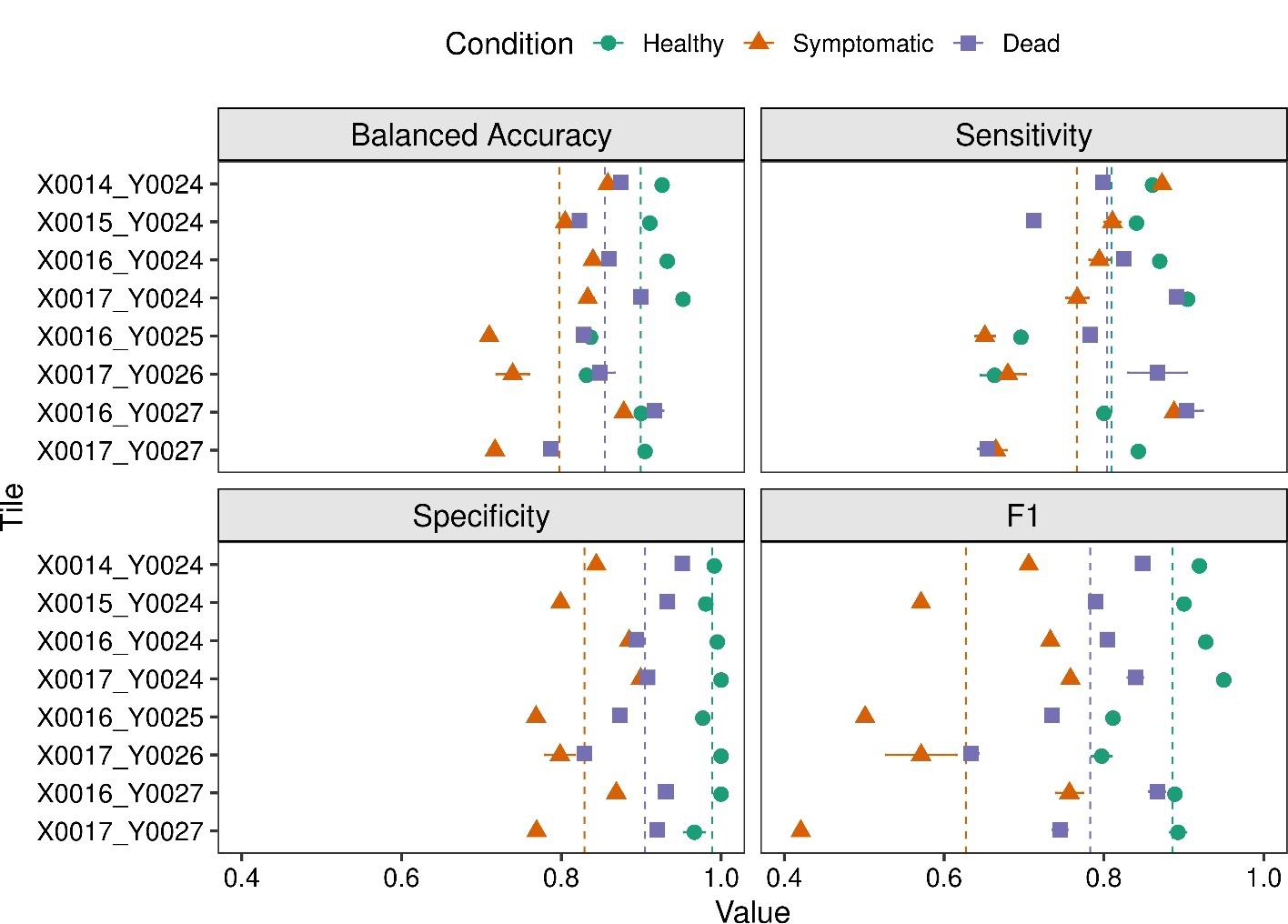
Spatial performance by tiles (900 km^2^) of the predicting model to discriminate between conditions of oak trees (healthy, symptomatic to oak wilt, and dead) on metrics derived from land surface phenology. Decreases in latitude (North-South) are described by increases in ‘Y’. Points of different shapes represent the mean, while the error bars are the standard deviation of the iterative models (*n* = 100). Vertical dotted lines describe the average per condition regardless of the tile. The spatial distribution of the tiles can be seen in Fig. S3.

#### 4.2.3 Temporal assessment

The temporal evaluation of the model built from 2019 data using the withheld 2019 data as well as the 2018 and 2021 datasets indicates that it is possible to classify oak tree conditions accurately through time (Fig. 7, Fig. 8). Similar to spatial performance, the temporal comparison reveals that pixels from healthy oak trees have higher values of balanced accuracy and F1 than pixels from symptomatic or dead oak trees (Fig. 7). Also, decreases in the sensitivity of pixels for symptomatic oak trees tend to be associated with sensitivity increases of pixels for dead and healthy oak trees (**Fig. S5**). The ROC curves (Fig. 8) and AUC (**Table S2**) also demonstrate that pixels of healthy oak trees are more accurately classified (AUC = 0.95 ± 0.02) than pixels for dead (AUC = 0.87 ± 0.02) or symptomatic oak trees (AUC = 0.79 ± 0.04). The macro (0.87 ± 0.02) and micro (0.88 ± 0.02) metrics of multiclass discrimination also indicate a congruent balance of classification performance independently of the conditions or class imbalance, respectively. Moreover, the probabilities of ‘*cutoff*’ to obtain a balance between sensitivity and 1 – specificity within conditions were relatively similar to the spatial assessment with average values of 0.42 (± 0.02), 0.29 (± 0.02), and 0.36 (± 0.01) for healthy, symptomatic, and dead oak trees respectively.

### 4.3 Mapping of oak tree conditions

The implementation of the model in areas outside of our training tiles enabled mapping symptomatic trees to oak wilt disease across locations (Fig. 9) and years (Fig. 10). For instance, pixels from symptomatic oak trees on forest patches (e.g., Fig. 9a) or outside (e.g., Fig. 9b) of pockets of disease tend to present contrasting probabilities that help to differentiate them among other pixels. In most of the cases, pixels that resample any of the evaluated conditions were also accompanied by lower values of overall uncertainties (e.g., Fig. 9 and Fig 10). In many instances edges of forest patches or areas that were not properly masked tend to be wrongly predicted as symptomatic or dead pixels (e.g., top right Fig. 9h); however, these pixels also tend to present high values of overall uncertainty. In this sense, the overall uncertainty appears to help isolate potential conditions (of any of the evaluated conditions) from false positives or other elements on the landscape such as grasslands (e.g., top right in Fig. 9c or **Fig S6j**), roads (e.g., top in Fig. 10h), or water bodies (e.g., left in Fig. 9f). Moreover, there were some instances where the predicted maps appeared with high values of overall uncertainty (Fig. 10g) and misleading probabilities (Fig. 10d and Fig. 6f), but this could be attributed to the reduced number of clear sky observations for that phenological year (e.g., **Fig. S2**).

**Fig 6.**
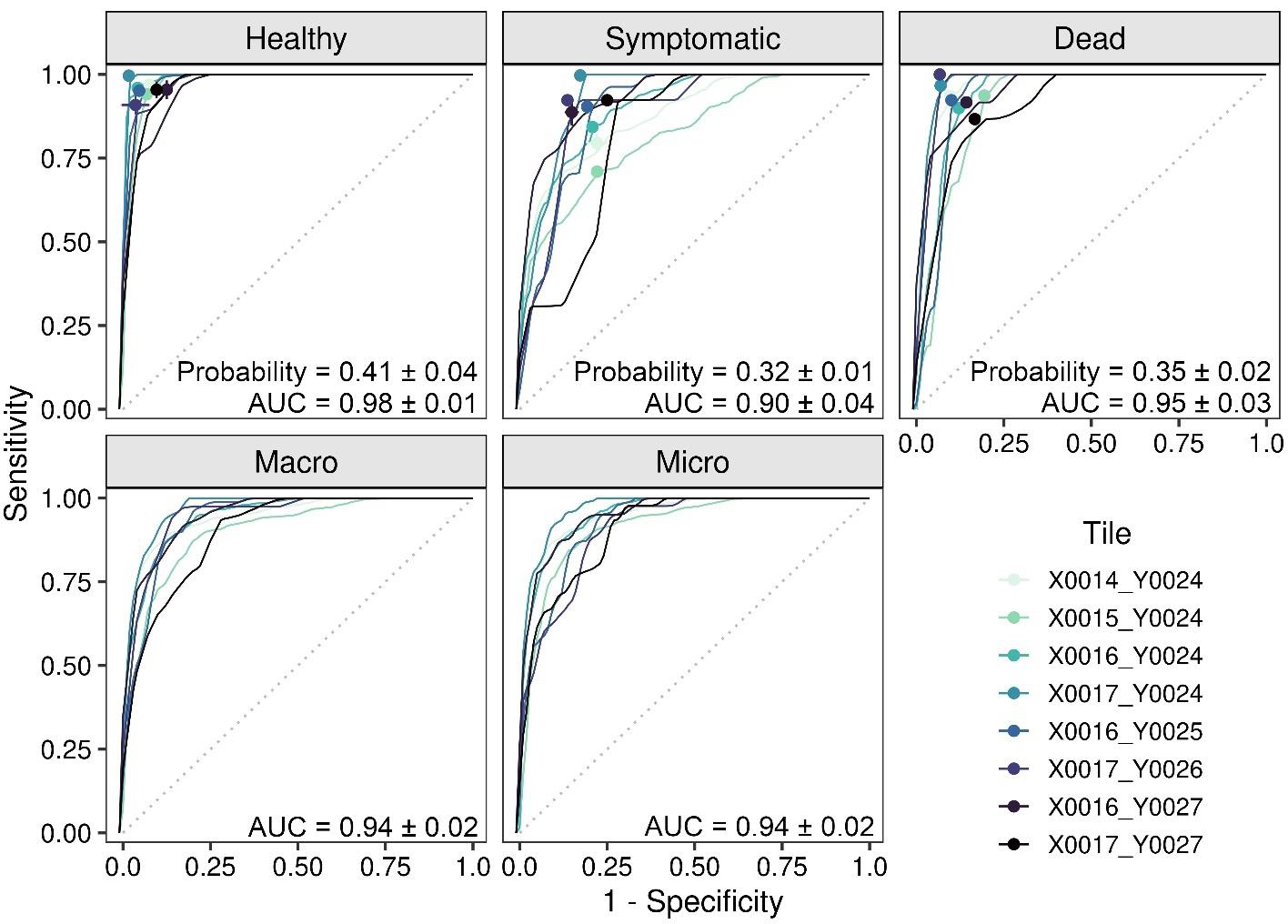
Spatial comparison of the Receiver Operating Characteristic (ROC) curves among tiles for each oak tree condition (healthy, symptomatic of oak wilt, and dead) and multiclass classification (macro and micro). Each line represents the mean of the iterative models (*n* = 100). The probability of cutoff among conditions and the Area Under the ROC Curve (AUC) describes the iterative models’ average (± SD) (*n* = 100) regardless of the tile. The gray dotted line represents the 1:1 relationship. The spatial distribution of the tiles can be seen in Fig. S3.

**Fig 7.**
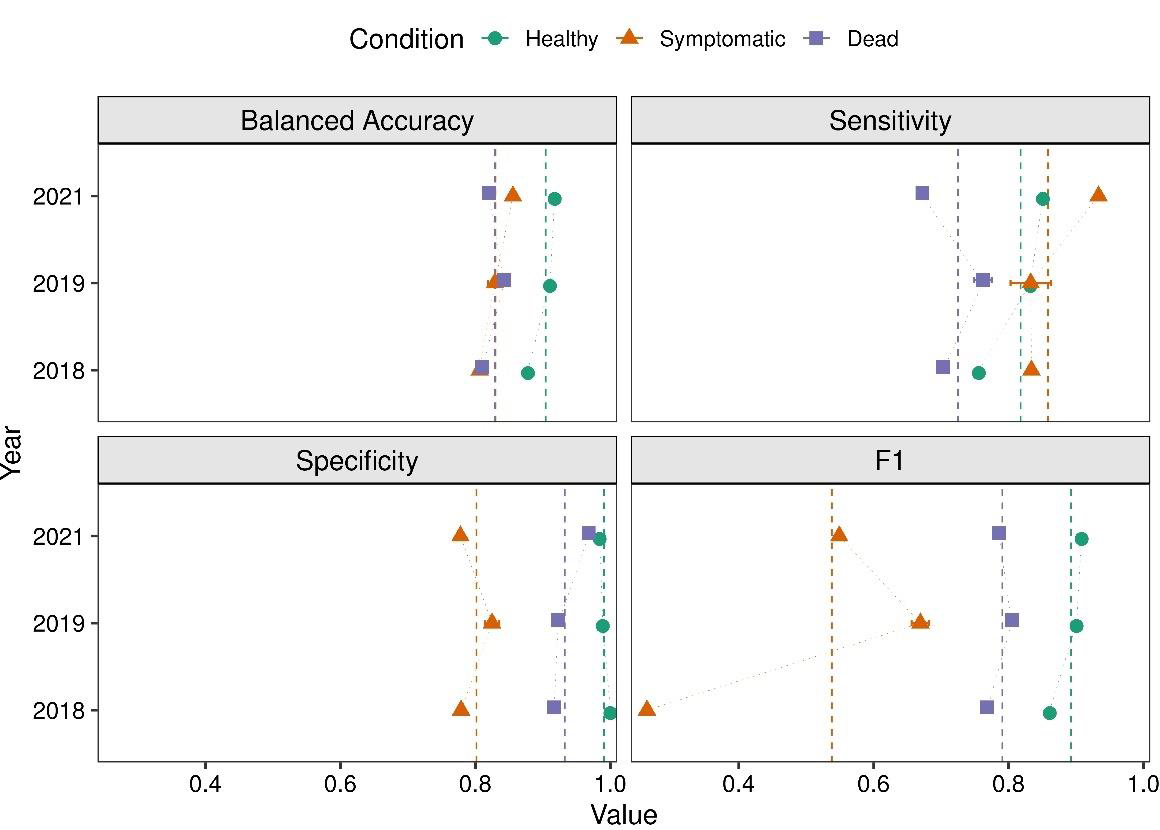
Temporal comparisons of model performance for predicting oak tree conditions (healthy, symptomatic of oak wilt, and dead) on metrics derived from land surface phenology. Points of different shapes represent the mean, and error bars are the standard deviations of the iterative models (*n* = 100). Vertical dotted lines describe the mean value per condition for all years.

**Fig 8.**
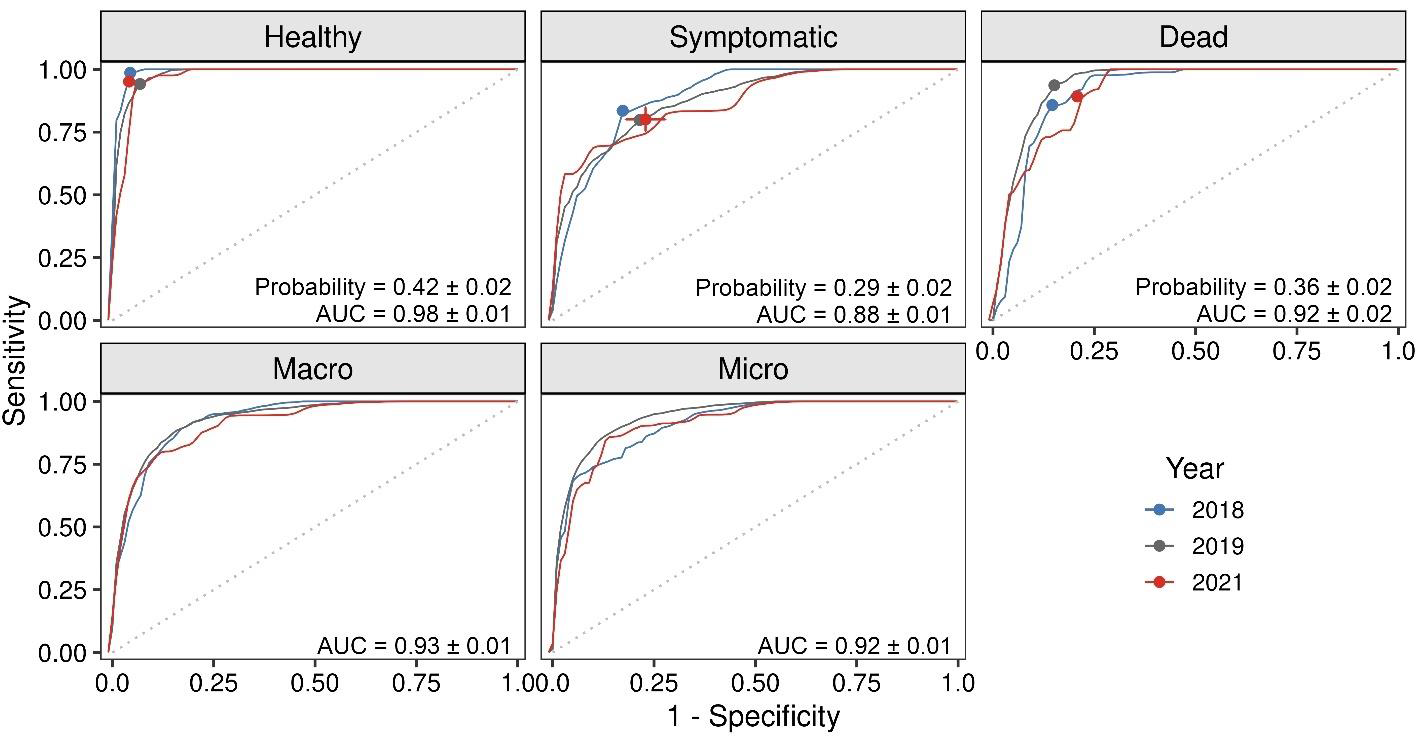
Temporal comparisons of the Receiver Operating Characteristic (ROC) curve across observations years for oak tree conditions (healthy, symptomatic of oak wilt, and dead) and multiclass classification (macro and micro). Each line represents the mean of the iterative models (*n* = 100). The probability of cutoff among conditions and the Area Under the ROC Curve (AUC) describes the iterative models’ average (± SD) (*n* = 100) regardless of the year. The gray dotted line represents the 1:1 relationship.

**Fig. 9.**
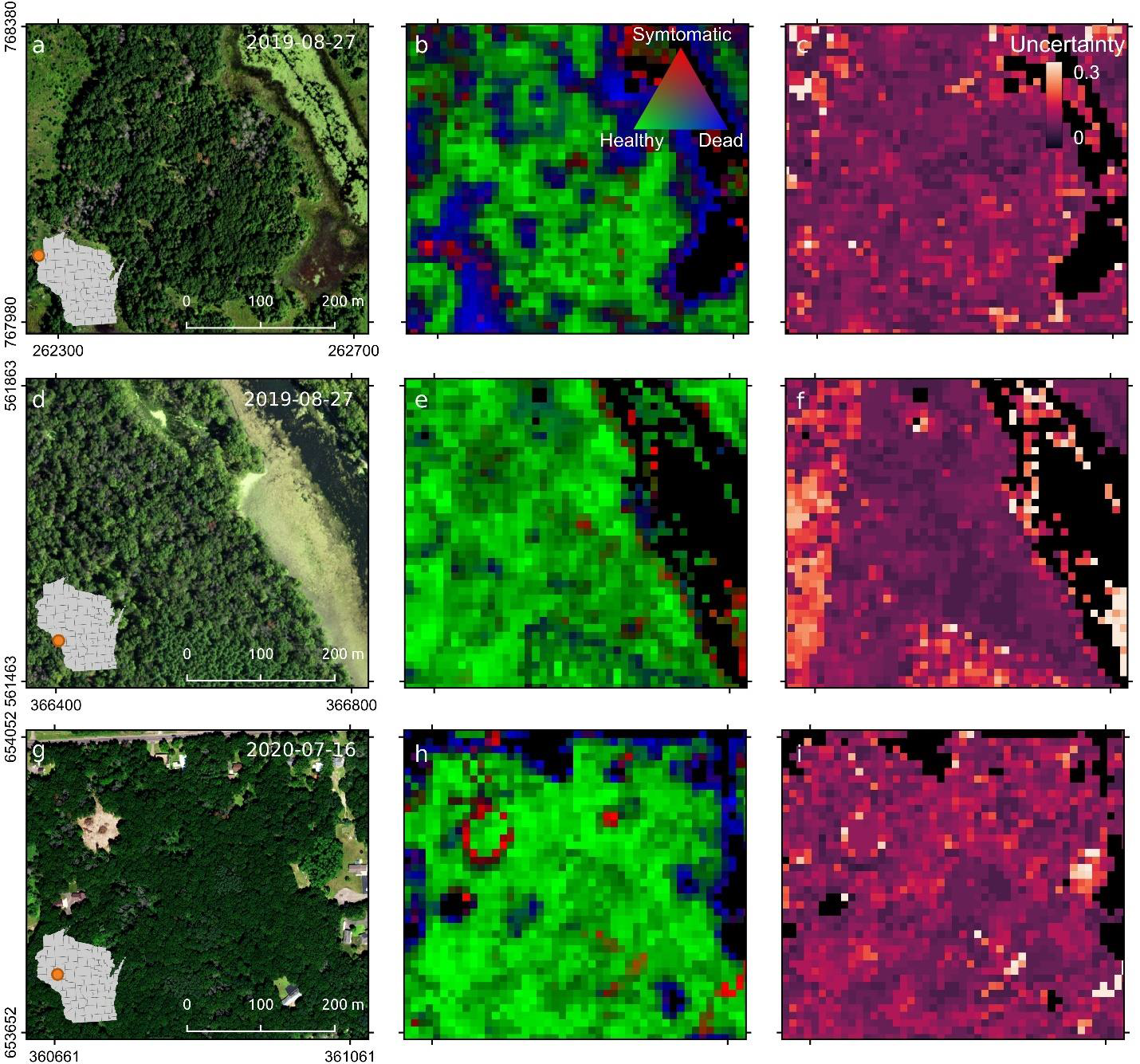
Mapping of oak wilt disease from healthy and dead oak trees across East Wisconsin. Panels **a**, **d**, and **g** are NAIP imagery from 2019 and 2020. Panels **b**, **e**, and **h** represent the predicted probability for healthy, symptomatic, and dead oak trees for the same years, while panels **c**, **f**, and **i** describe their overall uncertainty. The values of predicted probabilities were truncated between 0.3 and 0.6 for better visualization.

**Fig. 10.**
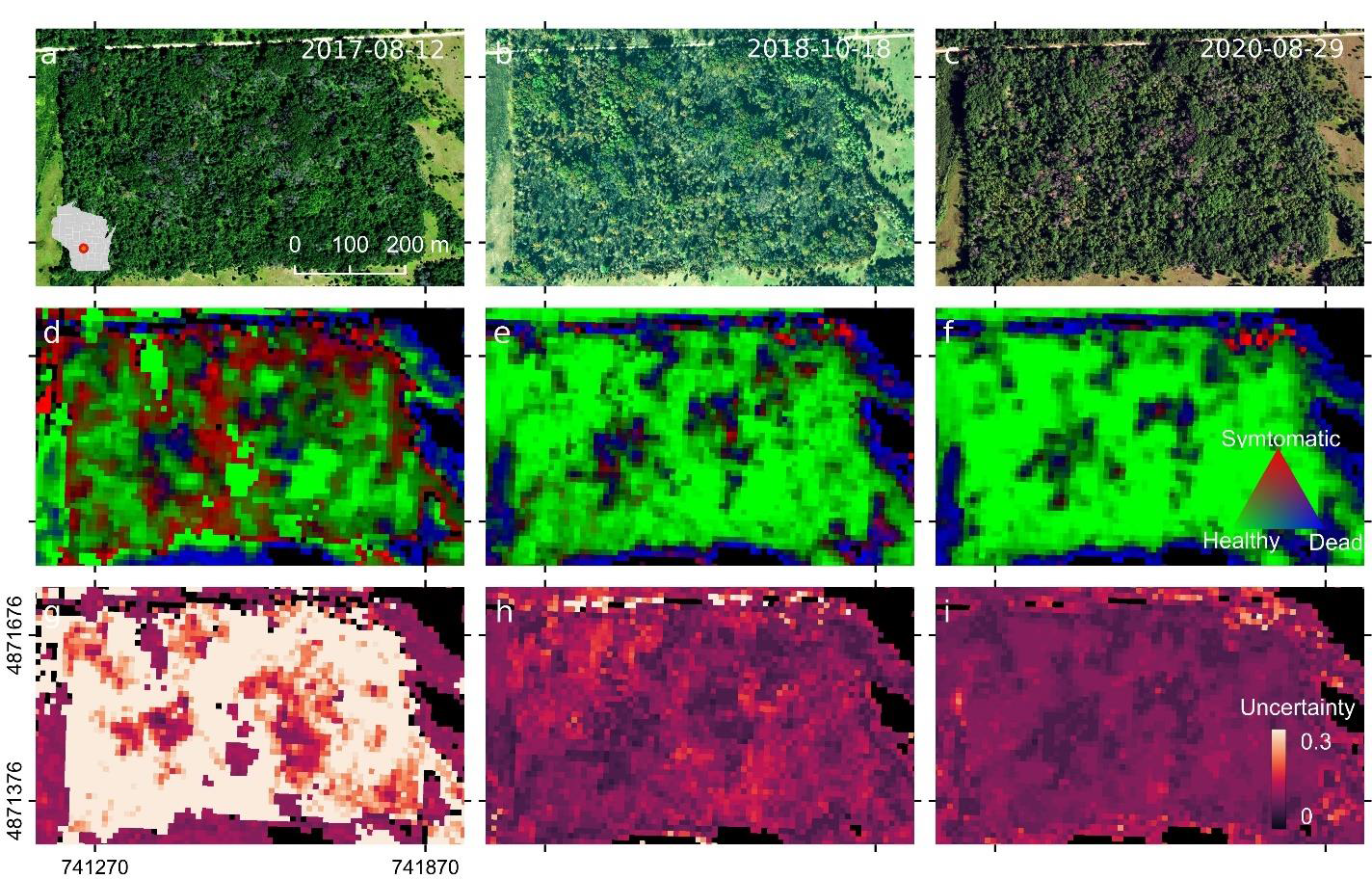
Temporal comparisons of the predicted probabilities of a pixel of being healthy, symptomatic of oak wilt disease, and dead in a forest patch at Buckhorn State Park, Wisconsin. Panels **a**, **b**, and **c** are NAIP images from 2017, 2018, and 2020. Panels **d**, **e**, and **f** represent the predicted probability for healthy, symptomatic, and dead oak trees for the same years, while panels **g**, **h**, and **i** describe their overall uncertainty. The values of the predicted probabilities were truncated between 0.3 and 0.6 for better visualization.

## 5. Discussion

Our results demonstrate a viable approach for classifying, mapping, and monitoring the presence of oak wilt using phenological observations from space. This approach targets changes in physiological symptoms of disease in oak trees using the chlorophyll carotenoid index (CCI) and examining the mechanistic basis for the temporal behavior of the oak trees with and without symptoms of oak wilt. We developed predictive machine learning models and assessed the approach spatially and temporally showing congruent and accurate classification of oak tree condition. We further unpack and highlight key aspects of the approach and offer insights into the future application of phenological observations to detect diseases at a large scale.

### 5.1 Spatial and temporal detection of oak wilt

Spatial and temporal evaluations showed that discrimination of symptomatic oak trees from healthy or dead trees reached values of accuracy comparable to those from airborne hyperspectral observations (∼ 80%) (Sapes et al., 2022). These discrimination results show that LSP metrics, which take advantage of physiological changes in tree canopies through time, are indirect and robust descriptors of disease. These descriptions appear to be coherent among classification algorithms during the training process, and robust spatially and temporally during the testing of the selected algorithm. The performance on testing datasets appeared to be slightly higher than training datasets which could be due to the balance sample selection from conditions during the training of the algorithms which is lacking in testing datasets. The iteration procedure selecting 80% of the samples during the training of the algorithms could also cause a large variability of performance on training datasets in comparison with testing datasets.

Although our results are promising for monitoring oak wilt, challenges associated with class confusion, local geography, edge effects, and data availability must be considered. First, symptomatic and dead oak trees are frequently confused. Overlapping values between conditions in the LPS metrics (i.e., Fig. 2 or **Fig. S3**) indicate that this likely results from spectral mixing, possibly combined with low temporal variability in signals of the disease symptoms. If an affected oak crown does not fully occupy a pixel the resulting signal will be mixed with surrounding elements (e.g., healthy or dead crowns). This effect is most likely to occur when only a single oak crown in a forest is affected (i.e., a crown is than a 10×10 m pixel or does not fully occupy a single pixel), or if the oak trees are small and their crowns are partially covered by other trees. This can also occur when oak wilt symptoms are discontinuous within a crown of a tree, as is common with white oaks. As such, our maps are likely more accurate for mapping oak wilt on red oaks than white oaks species; however, we have not explicitly tested this. In addition, the effect of spectral mixing also appears to be common at the edges of forest patches where tree crowns can mix their signals with other elements in the landscape. Observations within the interior of forest patches are therefore likely to have higher accuracy. Moreover, low temporal variability in signals of disease symptoms could be influenced by how the disease is expressed in the canopy or the number of observations through time, a function of image availability. For instance, few observations during the phenological cycle might result in poor differentiation of symptomatic oak trees from healthy or dead trees. Likewise, it is likely that the reduced number of available observations for 2017 (i.e., **Fig. S2**) makes predictions of that year more prone to high uncertainties (e.g., Fig 10g) compared to years with higher numbers of available observations. In addition, the spatial assessment showed that classification in three tiles had low performance compared to the average across all times for the three tree health conditions. This pattern may be attributable to the relatively few training samples that come from those tiles or the heterogeneous topography where these samples were collected (e.g., **Fig. S1**).

### 5.2 Mechanisms for detection of symptomatic trees to the disease

Three factors contribute to the accurate mapping of trees symptomatic of oak wilt in the approach we have presented: i) the sensitivity of the spectral index, ii) the phenological signal of the symptoms that are crucial to differentiation among oak tree conditions, and iii) the spatial resolution of the satellite imagery. The chlorophyll/carotenoid index (CCI) was used to target regions of the spectrum (i.e., red and green regions of the visible spectrum) that are associated with photochemical pigments that are influenced by the disease (Fallon et al., 2020). Although water absorption regions may also inform the presence of symptoms (Fallon et al., 2020; Sapes et al., 2022), changes in water stress on single tree crowns could be difficult to detect from spaceborne observations given both spectral contaminations in these regions with the atmospheric column and the typically coarse pixel size of bands at higher wavelengths. Other spectral indices have been tested to infer wilting-like diseases such as Normalized Difference Vegetation or Water Index (NDVI and NDWI, respectively) (De Castro et al., 2015; Kim et al., 2018; Sapes et al., 2022), and may be appropriate for other applications, but CCI likely performed best for oak wilt because of its direct relationship to the physiological effects of the disease. In general, the use of spectral indices (rather than all bands) to target spectral regions in conjunction with LSP metrics reduces data volume, which is critical to the feasibility of the workflow at a large scale. As well, spectral indices such as CCI from airborne sensors have also been shown to be quite promising for differentiating healthy trees from those with oak wilt in single time-point observations (Sapes et al., 2022).

Although it is essential to use a spectral index that is sensitive to a symptom, the phenological behavior of such a spectral index is critical for the discrimination among oak tree conditions. Seasonal changes in CCI have been associated with the regulation of photosynthetic activity via light harvesting and photoprotective processes (Springer et al., 2017; Wong et al., 2020). The CCI seasonality has been considered a reliable indicator of productivity in evergreen and broadleaf species (Gamon et al., 2016; Wong et al., 2020). In the context of oak wilt, where disease progression influences physiological processes across the season, metrics that capture temporal variation in CCI are powerful indicators of the disease (e.g., Fig. 1a). However, oak wilt may not be unique in its CCI temporal signal. For example, seasonal drought coupled with increased infestations by the Two-lined chestnut borer beetle (*Agrilus bilineatus*) could trigger phenological and physiological responses similar to oak wilt, resulting in an inability to distinguish between disease and climatic droughts or insect attacks. For the case of climatic droughts, however, the spatial pattern of drought-wilting is likely to be different from oak wilt disease. Climatic droughts tend to impact trees regionally, while the oak wilt mainly affects tree individuals or clumps of trees locally, often referred to as “oak wilt pockets”. On the other hand, wilting or mortality on oak trees due to outbreaks of the two-lined chestnut borer are also associated with reduced tree vigor due to drought, diseases, or defoliation (Haack and Acciavatti, 1992). Susceptible oak trees attacked by this beetle tend to wilt their leaves on scattered branches during late summer but keep them attached for several weeks or months before dropping (Haack and Acciavatti, 1992) in contrast to the early dropping of oak wilt symptomatic leaves. In addition, oak wilt-affected trees may also be infested with two-lined chestnut borer thus further complicating both “on-site” as well as remote detection. Attention to spatial and temporal differences in the impacts of regional drought or pest outbreaks is likely to help infer true positives for oak wilt.

Finally, spatial resolution is essential to detect disease on a single tree. This is relevant because the detection of single trees or pockets of disease is a critical need for forest managers looking to identify and contain outbreaks. A 10 m pixel size —as used in this research— can encapsulate or cover a proportion of a single crown, helping to reduce spectral mixing and thus providing a true wilting signal. However, it is more likely that in the case of a single-tree pocket, the crown will be distributed across two or more pixels, reducing the likelihood of detection (which can be further confounded by variation in pixel registration). Similarly, small oak trees will mix their signal with healthy or dead vegetation, hindering their detection. Other satellite imagery with higher spatial resolution such as those from the Dove constellation of PlanetScope or WorldView from Maxar can cover a proportion of tree crowns within a pixel with the potential to provide fine mapping of the disease. However, variable viewing geometry of these scenes, poorer signal-to-noise ratio, and the temporal harmonization of observations from different sensors might pose challenges to their application on a large scale (Teillet et al., 2007). Accessibility of this data (i.e., cost) may also pose issues for different stakeholders. Consequently, a coarse pixel size with better data quality could be more valuable than fine-resolution imagery for deriving LSP metrics (Helman, 2018), particularly if these data are readily available at high temporal frequency.

Our analyses focused on targeting and mapping symptomatic trees of oak wilt. However, because other diseases or disturbances may present temporal patterns in CCI similar to those that are symptomatic of oak wilt, identifying species or tree groups (lineages) will be a useful first step to diagnosing oak wilt. Differentiation among lineages of oak trees, for instance, is powerful for identifying oaks more susceptible to oak wilt (i.e., red oak species) using airborne hyperspectral imagery (Sapes et al., 2022). In comparison to high spatial resolution (<1 m) hyperspectral observations (Sapes et al., 2022), however, multispectral satellite observations (10 m) do not provide accurate lineage identity information sufficient to classify susceptible red oaks. Nevertheless, the discrimination of oak trees from other tree groups may help to isolate false positives of disease and prioritize areas with signs of new infection or areas of high infection density that pose risks of further spread. We emphasize that, to be useful, the predicted maps presented here still require expert knowledge from forest health specialists to validate symptomatic pixels to determine whether treatment is appropriate.

### 5.3 Future perspectives for disease detection using land surface phenology

Continuous satellite observations are valuable tools for assessing seasonal patterns. Deriving LSP metrics from such dense observations helps to characterize ecosystem function and the responses of forests to spatial and temporal stresses. Remote sensing of LSP has been extensively used in the literature to explore the mechanistic drivers of ecosystem phenology at spatial and temporal scales unfeasible in ground-based field surveys (Dronova and Taddeo, 2022). However, these kinds of approaches have rarely been applied to disease detection, despite their strong potential to advance forest pathogen monitoring.

Unlike detecting mortality, disease detection relies on the observation of visual (broad electromagnetic spectrum) symptoms (e.g., wilting, loss of leaves, reduction in pigments) that may enable the detection of the presence of a pathogen in advance of mortality. Monitoring and rapid detection at large spatial scales are critical first step to management efforts aimed at halting the spread of pathogens below- or above-ground. The temporal progression of symptoms of a disease is likely to be better represented by LSP metrics than a single or series of snapshots in time. LSP metrics leverage reductions in short-term signal variation (e.g., clouds, shadows, smoke) and data gaps imposed by single or bi-temporal observations (Scheffler and Frantz, 2022). Although the application of LSP metrics is slightly more computationally demanding, they can help to reduce the data volumes inherent in the use of continuous time series observations (e.g., average monthly observations as in Long et al. (2023)); speeding up the application of predictive models.

Dozens of phenological metrics that may inform a disease’s progression can be computed in a phenological cycle from spectral bands or indices (e.g., Jönsson and Eklundh, 2004). However, disease detection needs to be focused on bands/indices and metrics that are meaningfully connected to the symptoms to avoid artifacts that will hinder their discrimination and misinterpretation of the observed signals (Helman 2018). Therefore, efforts to decipher how disease progression and its changing symptoms over time drive the spectral properties in diseased vegetation are crucial and a first step for disease detection (Fallon et al., 2020). In addition, the selection of LSP metrics for detecting disease at large scales should also consider variation in phenology across latitudes. For instance, many LSP metrics may indicate the progression of oak wilt symptoms, including the cumulative value of CCI during the annual cycle or length of the growing period, but these metrics are likely to vary among latitudes (Bolton et al., 2020; Brooks et al., 2020).

We anticipate that the workflow and approach presented here could no doubt be leveraged for mapping other diseases and forest health at large scales, particularly on those diseases that show rapid progression within a single season. As such, future efforts focused on LSP metrics should include other pathogens and tree species.

## 6. Conclusion

This study demonstrates that it is feasible to detect and map the presence of oak wilt using phenological observations from space. Temporal changes in pigment composition caused by oak wilt fungus infection are key targets for discriminating symptomatic oak trees from healthy or dead trees using phenological metrics from satellite data. Discrimination is spatially and temporally coherent, indicating that it can be applied at large scales spanning multiple years. Challenges associated with spectral mixing, local geography, and data availability remain, but the approach and workflow provided will contribute to the efforts of stakeholders and managers in locating putative hotspots of disease, decreasing time to detection, and increasing management efficiency to slow the spread of this devastating tree disease. We anticipate that the approach presented here will leveraged for mapping other diseases at large scales, especially those with symptoms that can be tracked using vegetation indices. As high-frequency satellite-based CCI observations are becoming more readily available, phenological changes in disease-related physiology promise powerful applications for assessing tree diseases at large scales.

## Data and code availability

The code for processing the satellite data is available through https://github.com/davidfrantz/force, while the code for the z-score normalization of scenes and the development and prediction of the machine learning models is available at https://github.com/ASCEND-BII/Oak-wilt.git. The LSP metrics computed across states is available at https://app.globus.org/file-manager?origin_id=d5f9b461-7d6e-442b-87ed-be8aa2ca6763&path=%2F, while the predicted maps at https://app.globus.org/file-manager?origin_id=2ad70821-cc5a-424e-aa72-8553d2bb45eb&path=%2F.

## CRediT authorship contribution statement

**J. Antonio Guzmán Q.**: Conceptualization, Data curation, Formal analysis, Software, Investigation, Methodology, Visualization, Validation, Writing - original draft. **Jesús N. Pinto-Ledezma**: Writing - review & editing. **David Frantz**: Software, Writing - review & editing. **Philip A. Townsend**: Resources, Writing - review & editing. **Jennifer Juzwik**: Writing - review & editing. **Jeannine Cavender-Bares**: Conceptualization, Writing - review & editing, Resources, Supervision, Funding acquisition, Project administration.

## Acknowledgments

We thank the U.S. Geological Survey (USGS), U.S. Department of Agriculture, and European Space Agency (ESA) for providing free access to Landsat images, NAIP scenes, and the Copernicus Sentinel MSI for scientific research, respectively. This study was supported by the NASA Biodiversity Program (Award number: 80NSSC21K1349), the U.S. Forest Service (I20-CS-11330180-011), the Minnesota Invasive Terrestrial Pests and Pathogen Center, and the NSF ASCEND Biology Integration Institute (DBI: 2021898). We thank the UW2020 award from the Wisconsin Alumni Research Foundation (WARF) which supports the purchase of the HySpex instrument and Erin Wagner, Ben Spaier, W. Beckett Hills, Sonoma Brill, and Brendan Heberlein for the acquisition and processing of the imagery. Thanks to the Minnesota Supercomputing Institute for their high-performance infrastructure that allows us to conduct this research.

