## Supplementary material for "Mapping oak wilt disease using phenological observations from space": Table S1

**Table S1.** Total number of digitized points from healthy, symptomatic, and dead oak trees according to the year of observation and their spatial location in the data cube. The spatial distribution of these samples is represented in Fig. S3.

| Year of observation (Source) | Tile / Site | Condition | | | |
|---|---|---|---|---|---|
| | | Healthy | Symptomatic | Dead | Total |
| 2018 (HySpex) | X0014_Y0024 / Becker | 16 | 7 | 8 | **31** |
| | X0015_Y0024 / Cantlin | 25 | 6 | 11 | **42** |
| | X0016_Y0024 / Cedar Creek | 61 | 47 | 60 | **168** |
| | X0017_Y0024 / Stacy Dam | 6 | 4 | 5 | **15** |
| | **Total (2018)** | 108 | 63 | 84 | **256** |
| 2019 (NAIP) | X0014_Y0024 | 293 | 269 | 276 | **838** |
| | X0015_Y0024 | 380 | 351 | 368 | **1099** |
| | X0016_Y0024 | 309 | 265 | 288 | **862** |
| | X0017_Y0024 | 80 | 70 | 73 | **233** |
| | X0016_Y0025 | 71 | 65 | 64 | **200** |
| | X0017_Y0026 | 34 | 30 | 29 | **93** |
| | X0016_Y0027 | 32 | 28 | 28 | **88** |
| | X0017_Y0027 | 35 | 31 | 33 | **99** |
| | **Total (2019)** | **1234** | **1108** | **1159** | **3502** |
| 2021 (HySpex) | X0016_Y0024 / Cedar Creek | 41 | 36 | 37 | **114** |
| | **Total (2021)** | **41** | **36** | **37** | **114** |
| **Total** | | **1383** | **1209** | **1280** | **3872** |

**Table S2**. Mean Area Under the Curve (AUC) of the Receiver Operating Characteristic (ROC) curve of the iterative models to discriminate pixels from healthy, symptomatic, and dead oak tree conditions collected according to the year of observation and their spatial location in the data cube. These values correspond to the evaluations on the testing dataset. The standard deviation at all the levels of observation was lower than 0.02.

| **Year of observation (Source)** | **Level of observation (Tile / Year)** | **Condition** | | | | |
|---|---|---|---|---|---|---|
| | | **Healthy** | **Symptomatic** | **Dead** | **Macro** | **Micro** |
| 2018 (HySpex) | Year | 0.99 | 0.89 | 0.91 | 0.93 | 0.92 |
| 2019 (NAIP) | X0014_Y0024 | 0.99 | 0.88 | 0.97 | 0.95 | 0.95 |
| | X0015_Y0024 | 0.98 | 0.82 | 0.91 | 0.90 | 0.92 |
| | X0016_Y0024 | 0.99 | 0.91 | 0.94 | 0.95 | 0.95 |
| | X0016_Y0025 | 1.00 | 0.94 | 0.98 | 0.97 | 0.97 |
| | X0016_Y0027 | 0.99 | 0.90 | 0.94 | 0.94 | 0.93 |
| | X0017_Y0024 | 0.97 | 0.94 | 0.96 | 0.95 | 0.95 |
| | X0017_Y0026 | 0.99 | 0.90 | 0.98 | 0.95 | 0.91 |
| | X0017_Y0027 | 0.97 | 0.83 | 0.91 | 0.90 | 0.91 |
| | Year | 0.98 | 0.88 | 0.94 | 0.93 | 0.94 |
| 2021 (HySpex) | Year | 0.98 | 0.87 | 0.91 | 0.92 | 0.91 |
| **All testing observations** | | **0.98** | **0.89** | **0.94** | **0.94** | **0.93** |

**Table S3**. Required probability of obtaining a balance between true positives (Sensitivity) and false positives (1 - Specificity) derived from the Receiver Operating Characteristic (ROC) curve on iterative models to discriminate pixels from healthy, symptomatic, and dead oak tree conditions. Probabilities are described according to the year of observation and the spatial location in the data cube. These values correspond to the evaluations on the testing dataset. The standard deviation at all the levels of observation was lower than 0.02.

| **Year of observation (Source)** | **Level of observation (Tile / Year)** | **Condition** | | |
|---|---|---|---|---|
| | | **Healthy** | **Symptomatic** | **Dead** |
| 2018 (HySpex) | Year | 0.44 | 0.27 | 0.38 |
| | X0014_Y0024 | 0.38 | 0.32 | 0.36 |
| | X0015_Y0024 | 0.38 | 0.32 | 0.35 |
| | X0016_Y0024 | 0.41 | 0.32 | 0.34 |
| | X0017_Y0024 | 0.42 | 0.32 | 0.35 |
| 2019 (NAIP) | X0016_Y0025 | 0.44 | 0.32 | 0.35 |
| | X0017_Y0026 | 0.41 | 0.35 | 0.31 |
| | X0016_Y0027 | 0.50 | 0.33 | 0.33 |
| | X0017_Y0027 | 0.37 | 0.32 | 0.38 |
| | Year | 0.39 | 0.32 | 0.34 |
| 2021 (HySpex) | Year | 0.42 | 0.29 | 0.38 |
| **All testing observations** | | **0.41** | **0.32** | **0.35** |

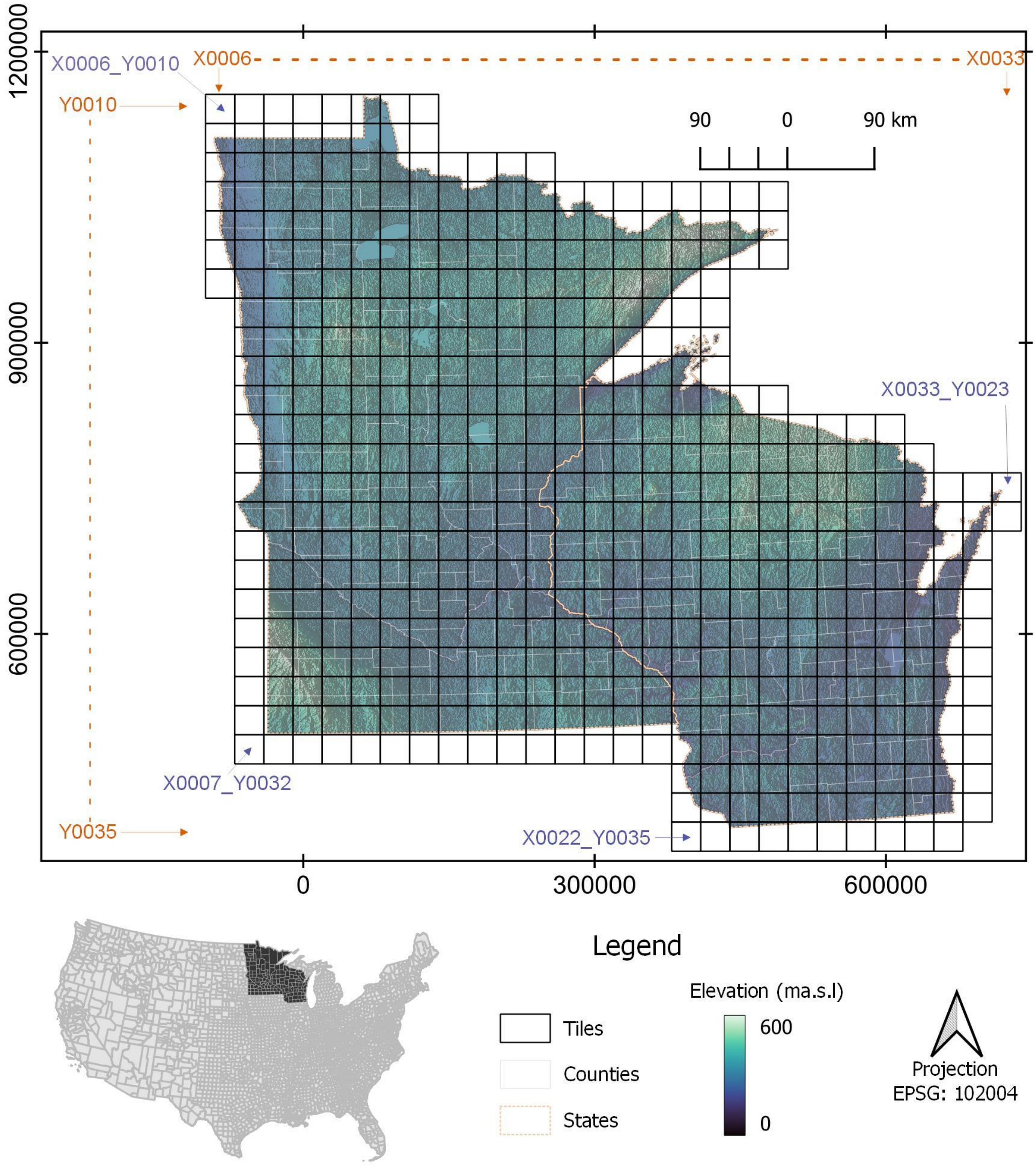


**Fig. S1.** Spatial distribution of the tiles in the datacube across Minnesota and Wisconsin.

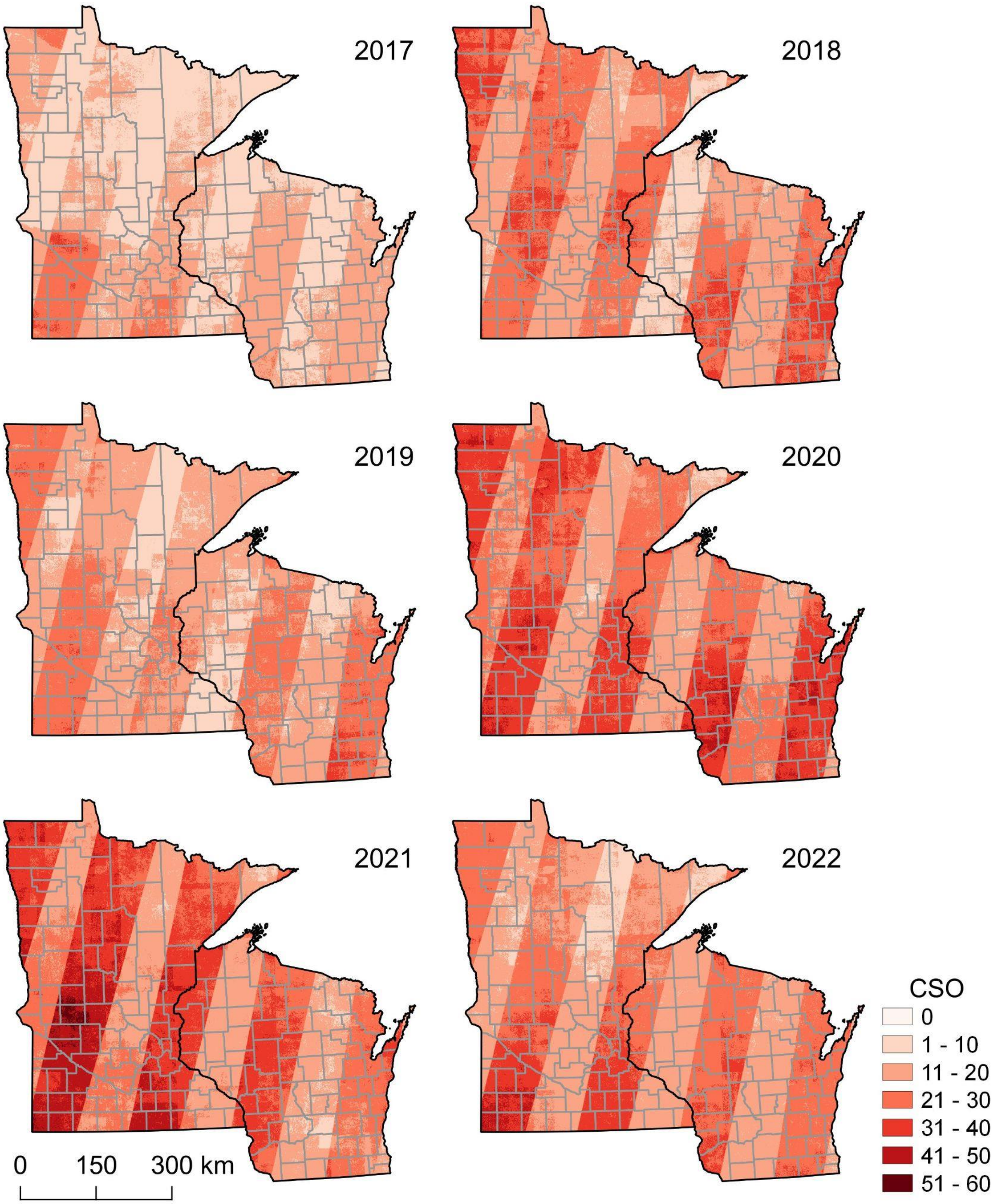


**Fig. S2.** Clear sky observations (CSO) of Sentinel-2 across Minnesota and Wisconsin from 2017 to 2022. The stripes describe a higher data availability due to the orbit overlap. Gray boundaries are counties.

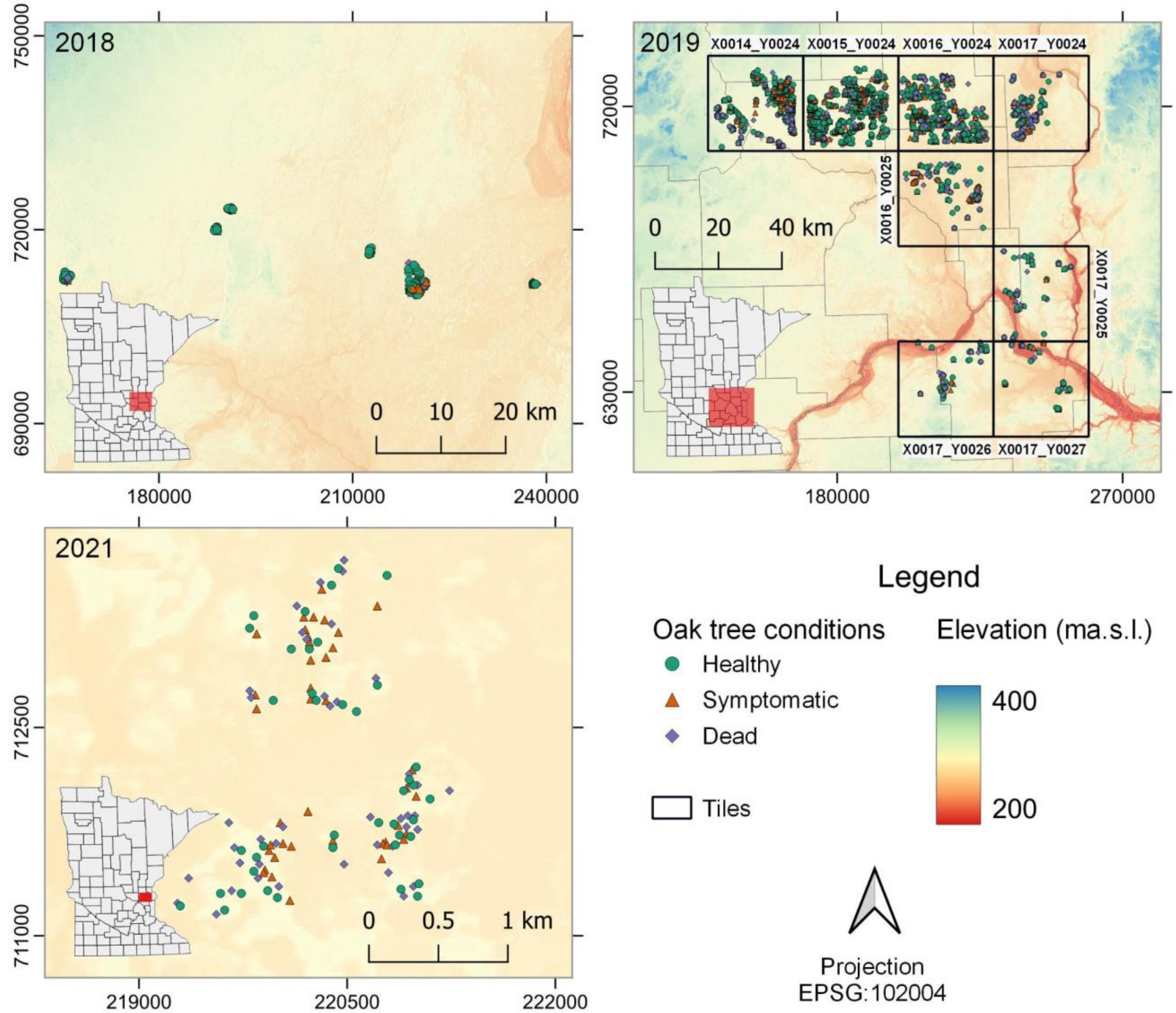


**Fig. S3.** Spatial distribution of the digitized points from healthy, symptomatic, and dead oak trees according to the year of observation and their location in the data cube.

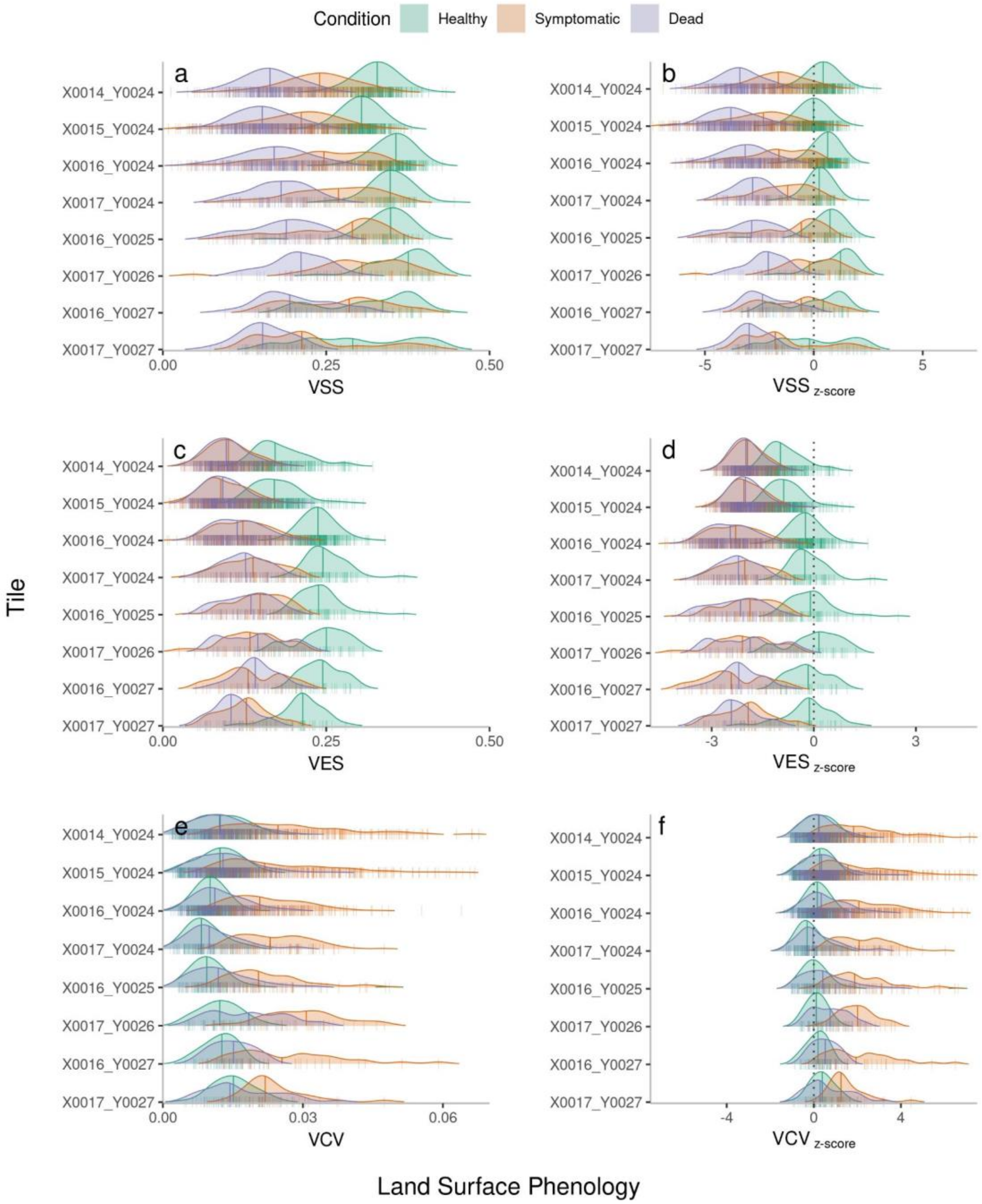


**Fig. S4.** Ridge plots of land surface phenology metrics (a, c, e) comparing three oak tree conditions among tiles and their z-score normalization (b, d, f). Value at the start of the Season (VSS) (a, b), Value at the End of the Season (VES) (c, d), and value of the coefficient of variation (VCV) (e, f). The irregular polygons describe the kernel density distributions with different bandwidth for better visualization. Each small vertical line represents a pixel (i.e., an oak tree crown), while the large vertical lines within the polygons the 50$^{th}$ percentile.

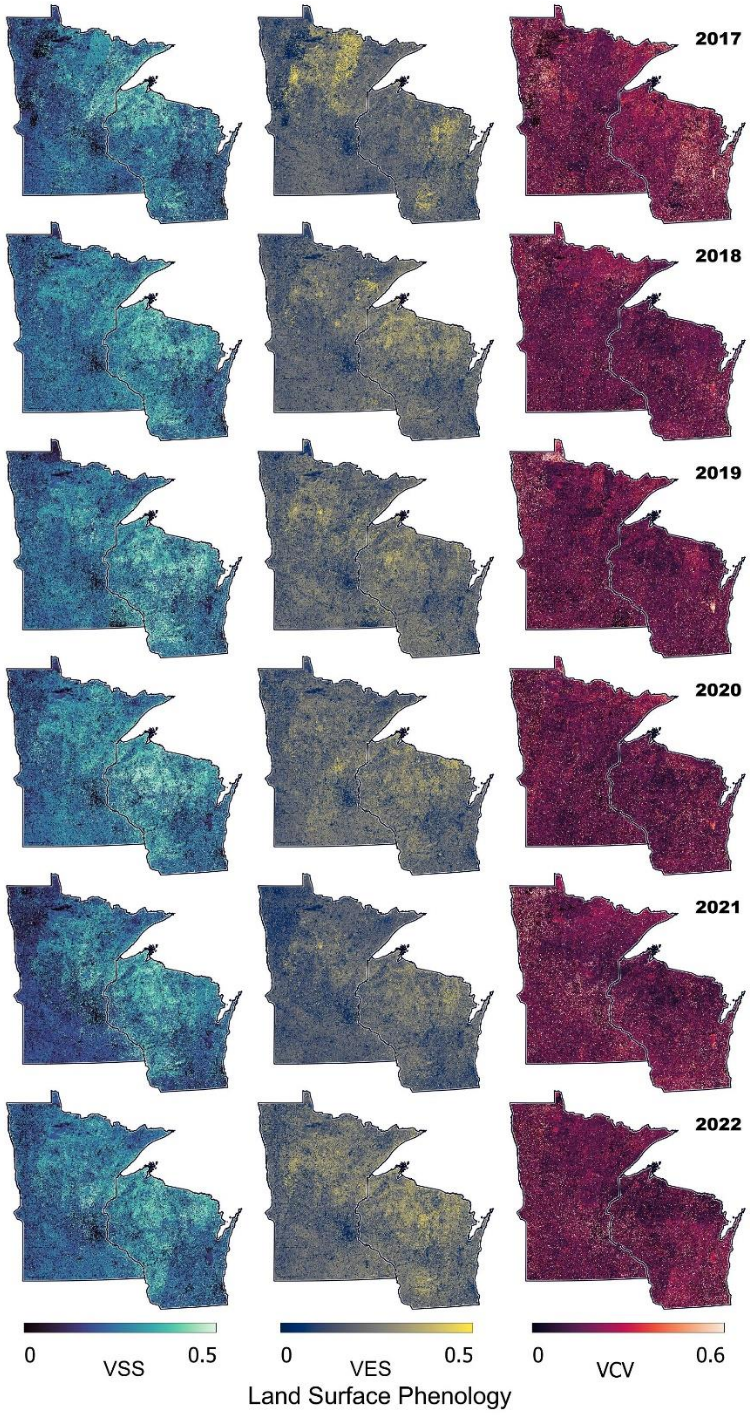


**Fig. S5.** Land Surface Phenology of the Chlorophyll/Carotenoid Index across Minnesota and Wisconsin derived from Sentinel-2 between 2017 to 2022. Value of the Start of the Season (VSS), Value at the End of the Season (VES), and Value of Coefficient of Variation (VCV) during the growing period.

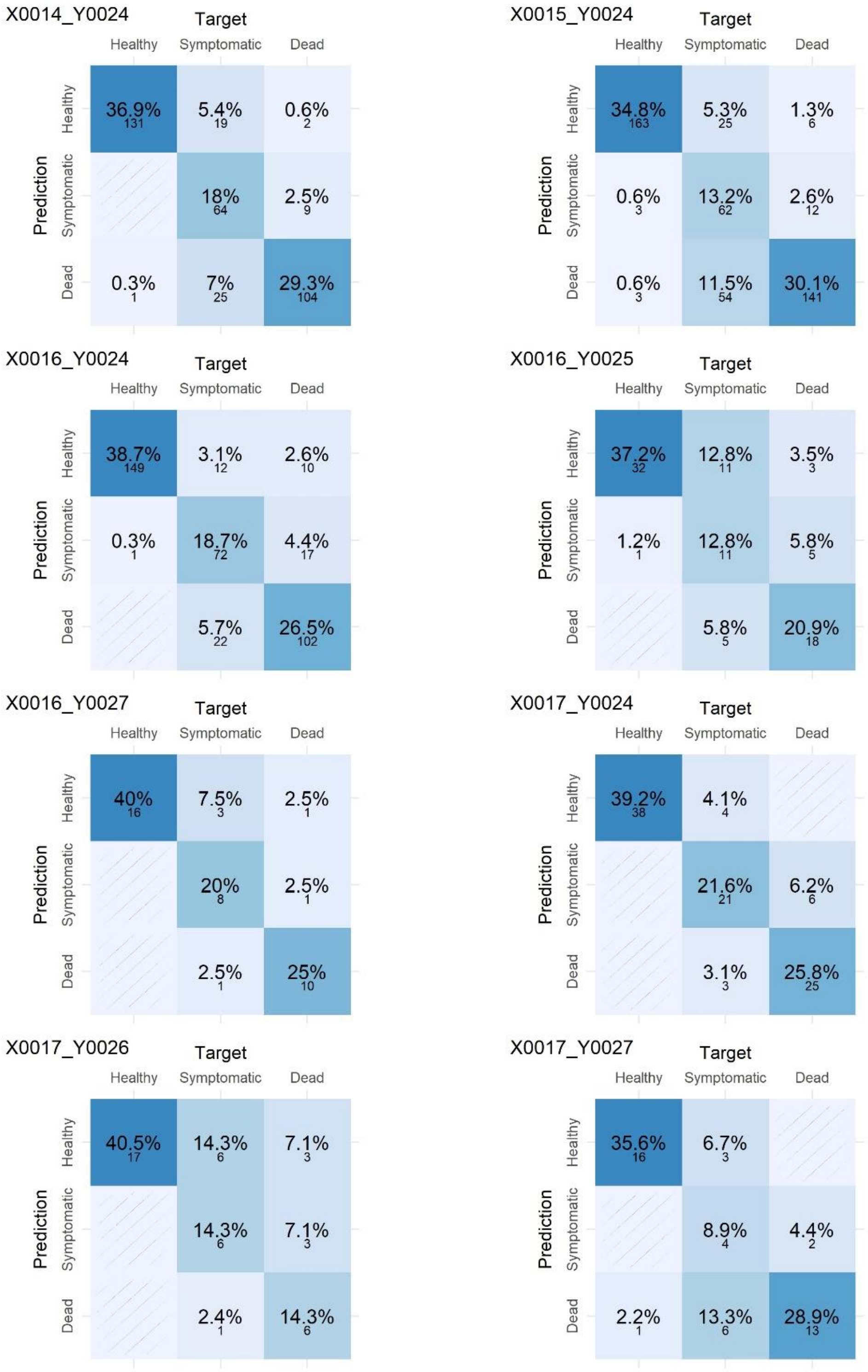


**Fig. S6.** Confusion matrices for the spatial assessment of the predicting model to discriminate between conditions of oak trees (healthy, symptomatic to oak wilt, and dead) on metrics derived from land surface phenology.

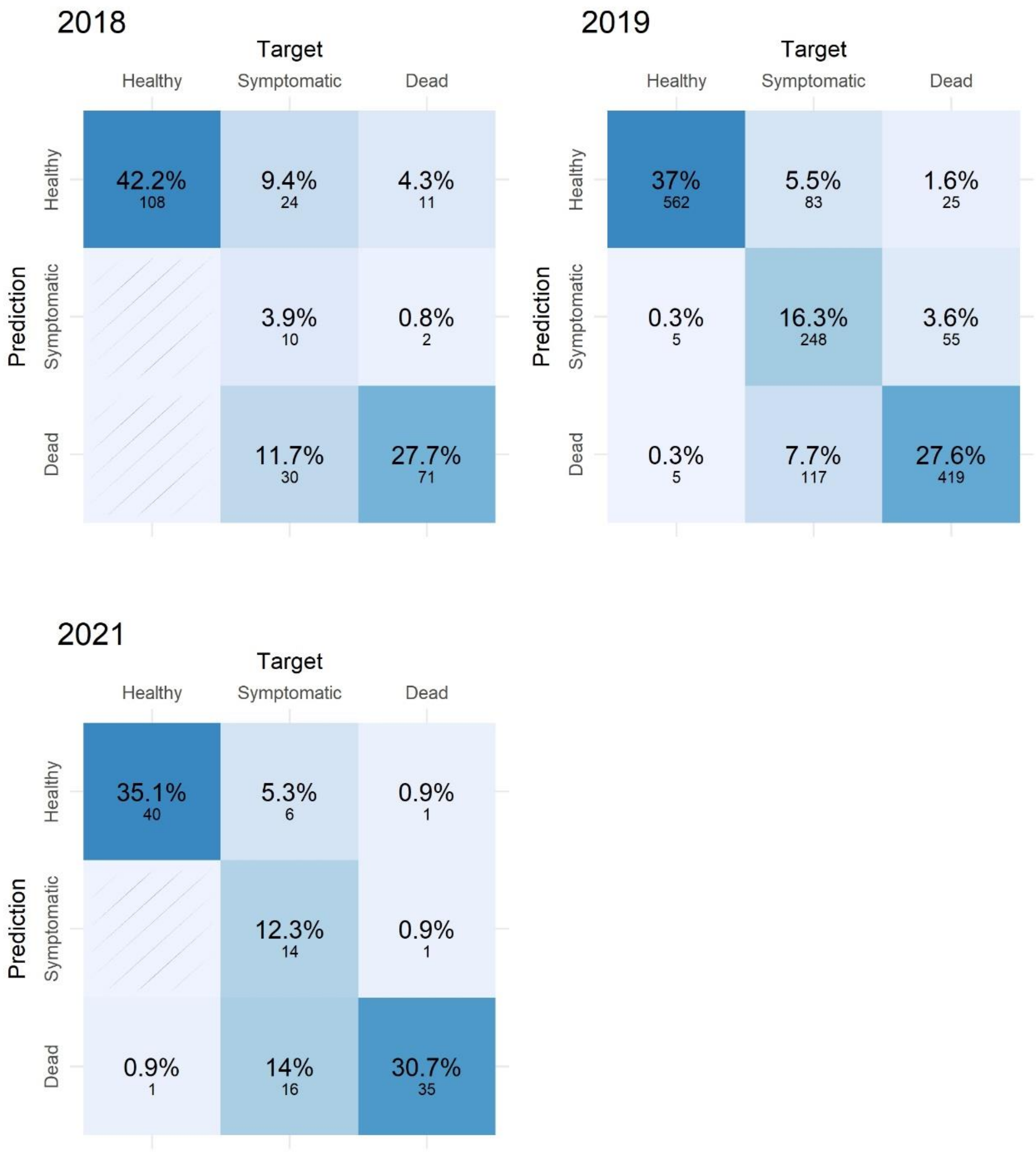


**Fig. S7.** Confusion matrices for the temporal assessment of the predicting model to discriminate between conditions of oak trees (healthy, symptomatic to oak wilt, and dead) on metrics derived from land surface phenology.

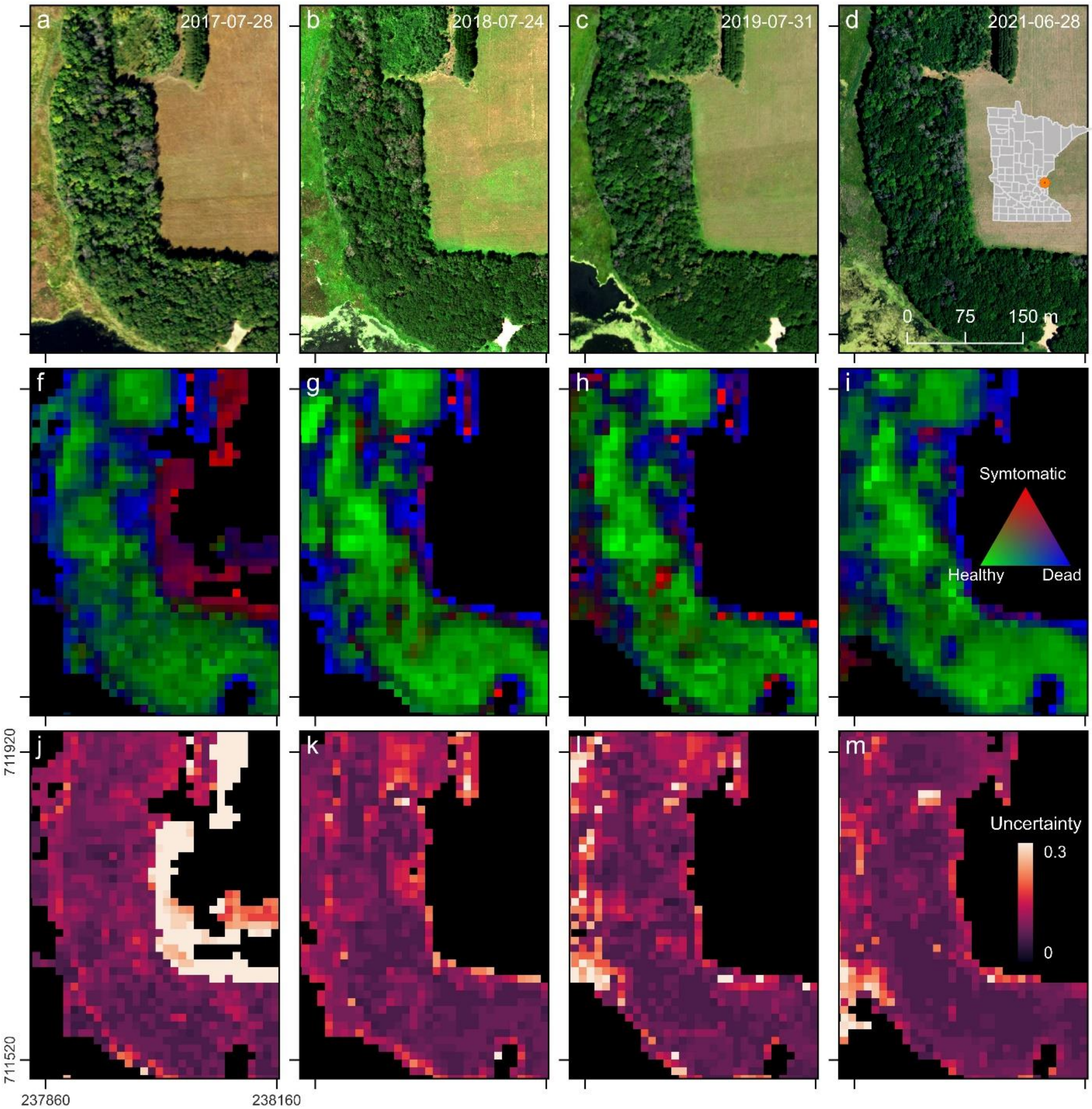


**Fig. S8.** Temporal comparisons of the predicted probabilities of a pixel of being healthy, symptomatic of oak wilt disease, and dead in a forest patch at Stacy Dam, Minnesota. Panels **a**, **c**, **b**, and **d** are NAIP and HySpex images from 2017, 2018, 2019, and 2021. Panels **f**, **g**, **h**, and **i** represent the predicted probability for healthy, symptomatic, and dead oak trees for the same years, while panels **j**, **k**, **l**, and **m** describe their overall uncertainty. The values of the predicted probabilities were truncated between 0.3 and 0.6 for better visualization.
